# The hybrid yeast *Saccharomyces pastorianus* TUM 34/70 repurposes the pseudohyphal growth signaling network to establish a flocculation phenotype

**DOI:** 10.64898/2026.08.03.742438

**Authors:** Katharina Seibel, Eoin Ó Cinnéide, Riley Schmalhaus, Mirjam Haensel, Florian Weiland

## Abstract

Lager is the most produced beer style world-wide and makes use of the bottom-fermenting hybrid yeast *Saccharomyces pastorianus* (*S. cerevisiae* x *S. eubayanus*). Previous research showed that flocculation in *S. pastorianus,* in contrast to the top-fermenting ale yeast *S. cerevisiae,* is triggered by nitrogen starvation. However, the cellular events leading to flocculation in *S. pastorianus* are not well characterized. Therefore, we conducted a proteomic screen of *S. pastorianus* TUM 34/70 and identified the protein kinase Ste20p and protein phosphatase regulatory subunit Ypi1p as higher abundant during flocculation. Overexpression of these genes caused a consistent and strong increase in flocculation rate over the complete duration of beer fermentation. Characterization of Ste20p and Ypi1p via a phospho-proteomics screen showed their targeting of proteins whose *S. cerevisiae* orthologues are involved in pseudohyphal growth. However, in contrast to this, the overexpression of *STE20* and *YPI1* led instead to the establishment of a flocculation morphology, giving first-time evidence that *S. pastorianus* repurposes the pseudohyphal signaling network for this phenotype.

## 1 Introduction

The brewing industry is an important economic factor world-wide, generating nearly 900 billion USD and supporting more than 33 million jobs in 2023 (Mjikane, 2025). The most produced style is lager with around 70% of the total 1.95 billion hl beer produced in 2025 (*Lager Market Size, Share | Industry Report, 2035*, n.d.) and for this beer type the bottom-fermenting brewing yeast *Saccharomyces pastorianus* is utilized.

*S. pastorianus* is an interspecific hybrid of *S. cerevisiae* and *S. eubayanus*, dating back to single hybridization event around 500 years ago (Salazar et al., 2019). Two subtypes within this species are known, Saaz and Frohberg, which are characterized by differing chromosomal re-arrangements (reviewed in (Wendland, 2014)), resulting in different cold resistances and sugar utilization capabilities (Gibson et al., 2013). As such, Frohberg yeasts are generally preferred in the brewing industry as they can utilize maltotriose, the second highest abundant sugar typically encountered in beer wort (Magalhães et al., 2016; Stewart, 2016) and the Frohberg strain TUM 34/70 is one of the most widely used in the brewing industry (Müller-Auffermann et al., 2015). A distinct feature of *S. pastorianus* is its flocculation behavior, in which the yeast cells clump together and sink to the bottom of the vessel at the end of beer fermentation. In the top fermenting *S. cerevisiae* flocculation results from the expression of the gene *FLO1*. The lectin Flo1p binds adjacent (phospho)mannan residues in the cell wall of other yeast cells, in turn leading to flocculation (reviewed in (Seibel et al., 2021; Soares, 2011; Verstrepen et al., 2003). In *S. pastorianus* the *FLO1* homologue Lg-*FLO1* is found and the protein Lg-Flo1 shares around 60% of its amino acid sequence with Flo1p (Kobayashi et al., 1998). Importantly, Lg-*FLO1* is translocated to the telomere of chromosome VIII and under control of a promotor resembling that of *FLO5* (Ogata et al., 2008) and flocculation in *S. pastorianus* is triggered by nitrogen starvation, which is in contrast to *S. cerevisiae* where flocculation is induced by lack of carbon source (Ogata, 2012).

Flocculation is of great importance to the brewing process, as it is low cost and environmentally friendly, enabling easy and efficient separation of yeast cells from beer. It is therefore warranted that flocculation in *S. pastorianus* is well understood, but the underlying cellular events remain largely unknown. Therefore, we analyzed the proteome of *S. pastorianus* TUM 34/70 during flocculation to gain insights into potentially involved cell signaling proteins. We identified the kinase Ste20p and phosphatase subunit Ypi1p as higher abundant in flocculent cells and validated their involvement in flocculation. Phosphoproteomic characterization of these proteins strongly suggest that the pseudohyphal growth pathway is repurposed in *S. pastorianus* TUM 34/70 to lead to flocculation instead.

## 2 Material and Methods

### 2.1 Media preparation

Growth medium for nitrogen source selection was prepared using un-hopped malt extract (Weyermann^®^ Bavarian Pilsner, Weyermann Specialty Malting, Bamberg, Germany). Malt extract was added into 60 °C deionized water until 12.8 °Plato (P) was reached as measured by an EasyDens density meter (Anton Paar GmbH, Graz, Austria) (measured at 20 °C). Resulting wort was cooled down to 9 °C and the fermentation was started immediately (see *2.4 Influence of nitrogen source on yeast flocculation behavior)*.

For yeast propagation and laboratory-scale beer fermentation, 12 °P wort was prepared by adding 745 g Muntons Spraymalt Light (Muntons Ingredients, Stowmarket, United Kingdom) to 3,544 g deionized water, the mixture was stirred and afterwards autoclaved at 121 °C for 20 min (LABOKLAV 135, SHP Steriltechnik AG, Schloss Detzel, Germany), cooled to room-temperature (RT) and sterile filtered with a 0.22 µm bottle top filter (Cat. No.: FB12566508, Thermo Fisher Scientific, Waltham, Massachusetts, USA) to remove undissolved particles. Sterile wort was stored at room temperature (RT).

YPS medium agar plates were prepared as follows: 100 g/L saccharose (Everyday, Halle, Belgium), 10 g/L yeast extract (Cat. No.: LP0021, Thermo Fisher Scientific), 5 g/L peptone (Cat. No.: LP0034, Thermo Fisher Scientific), 1.2 g/L ammonium sulphate (Cat. No.: 101217, Merck KGaA, Darmstadt, Germany), 1 g/L potassium dihydrogen phosphate (Cat. No.: 104873, Merck KGaA), 0.7 g/L magnesium chloride (Cat. No.: 197530010, Acros Organics, Geel, Belgium), 0.5 g/L sodium chloride (Everyday), 0.1 g/L calcium chloride (Cat. No.: 207780010, Acros Organics), 5 mg/L zinc chloride (Cat. No: AC198945000, Acros Organics) and 20 g/L agar (Cat. No.: L13, Unipath LTD, Basingstroke, Hampshire, England) were dissolved in 1 L deionized water, autoclaved at 121 °C for 20 min, poured into 90 mm plastic dishes (Cat. No.: 391-0601, VWR Avantor Inc., Randor, Pennsylvania, USA) and stored at 4 °C until usage.

For experiments regarding yeast transformation yeast extract, peptone, dextrose media (YPD) was used. For this. 10 g/L Yeast extract (Cat. No.: ACRO451120010, VWR Avantor Inc.), 20 g/L Bacto™ Peptone (Cat. No.: 211677, Thermo Fisher Scientific), 20 g/L alpha-D-Glucose (Cat. No.: 158968, Merck KGaA) was dissolved in 1 L ultrapure water, autoclaved at 121 °C for 20 min and stored at RT until further usage. For YPD agar plates, YPD medium was prepared as above with addition of 20 g/L agar (Unipath LTD). After autoclaving, the medium was cooled to approx. 45° C and Geneticin disulphate G418 (Cat. No.; CP11.2, Carl Roth GmbH & Co. KG, Karlsruhe, Germany) was added to a final concentration of 10 µg/mL.

### 2.2 Yeast cryoculture preparation

*Saccharomyces pastorianus* TUM 34/70 yeast was purchased at the Yeast Centre of the TU Munich-Weihenstephan on slanted agar (Frisingia TUM 34/70 yeast, Cat. No.: 95000a, TUM School of Life Sciences, Freising, Germany). The yeast was streaked out on a YPS-agar plate and incubated at 12 °C until colonies were visible (2-3 days). For preparation of cryo-stocks a colony was picked from the plate, inoculated in 10 mL wort and grown for 48 h at 12 °C before being mixed with 5 mL glycerol (final glycerol concentration: 33% (v/v), Cat. No.: G6279-500ML, Merck KGaA). The cryocultures were stored at-80 °C until further usage.

### 2.3 Yeast propagation

First, a cryoculture of *S. pastorianus* TUM 34/70 was streaked out on a YPS agar plate and incubated at 22°C until colonies were visible (around 5 days). From this plate, a smear of yeast was inoculated in 10 mL sterile 12 °P wort and incubated at 12 °C (n = 5). After inoculation for 48 h another 100 mL sterile 12 °P wort was added, and incubation was continued at 12 °C while stirring at 130 rpm. Upon reaching 96 h after start of inoculation an additional 500 mL sterile 12 °P wort was added and yeast propagation continued at 12 °C with stirring at 130 rpm for another 72 h (168 h total).

### 2.4 Influence of nitrogen source on yeast flocculation behavior

Three 5 L glass bottle containing each a magnetic stirrer and 1800 mL media were inoculated with freshly propagated 15 x 10^6^ cells/mL *S. pastorianus* TUM 34/70 at 9 °C with stirring at 130 rpm for 24 h. Afterwards, the yeast cultures were split into 250 mL and the different nitrogen sources were added respectively as solid powder: 1) 10 g/L ammonium sulphate (n = 3, Cat. No.: SAFSA4418-100G, VWR), 2) 1 g/L glutamine (n = 3, Cat. No.: SAFSG8540-25G, VWR) and 1 g/L glutamic acid (n = 3, Cat. No.: SAFA49621-250G, VWR), 3) 1 g/L proline (n = 3, Cat. No.: SAFSP5607-25G, VWR) or 4) 0.6 g/L urea (n = 3, Cat. No.: SAFSU5378-100G, VWR), while the negative control was left untreated.

Extract, cell density, pH and yeast flocculation rate were measured 24 h, 32 h, 48 h, 72 h and 96 h after inoculation. The flocculation rate was determined via Helm’s test according to Smart et al. (D’Hautcourt & Smart, 1999). Cell density was established by measuring the optical density at 600 nm (OD6_00_) (Novaspec II Spectrophotometer, Pharmacia, Uppsala, Sweden) and conversion into cells per mL via a calibration curve derived from a Thoma counting chamber (see **Supplementary Figure 1**). For manual counting, the cells were diluted 1:8 (v/v) with 0.05 M EDTA (Titriplex® III, Cat. No.: 1084311000, Merck KGaA). For extract and pH measurements, samples were filtered using a folded filter (Cat. No.: 6000508, Cytiva Europe GmbH, Freiburg im Breisgau, Germany) and Becogur 200 (Eaton Electric GmbH, Bonn, Germany). Samples were afterwards centrifuged for 5 min at 3,500 rcf at RT. The extract was determined using an EasyDens density meter (Anton Paar GmbH). The pH of the supernatant was measured using a pH electrode (Hanna Instruments Deutschland GmbH, Vöhringen, Germany).

### 2.5 Beer fermentation

Freshly propagated *S. pastorianus* TUM 34/70 were inoculated with a concentration of 15 x 10^6^ cells/mL in 1 L glass bottles containing each 1 L of sterile 12 °P wort. Samples were cultured at 12 °C and stirred at 200 rpm. 48 h after inoculation, yeast cultures (n = 5) were supplemented with 1 g/L L-glutamine (Cat. No. HN08.3, Carl Roth GmbH & Co. KG) and 1 g/L L-glutamic acid (Cat. No. 1743.2, Carl Roth GmbH & Co. KG), by weighing in the supplements directly into the culture, while control samples (n = 5) remained untreated.

Cell concentration of the overall culture and (after 15 minutes unstirred rest) of the sedimented and non-sedimented cells, was determined by measurement of OD_600_ (Agilent Cary 100 UV VIS spectrophotometer, Agilent, Santa Clara, United States) and converted to number of cells per mL with an in-house generated calibration curve (**Supplementary Figure 2**). The calibration curve was constructed by plotting the measured the OD_600_ value over the corresponding cell density determined by counting the cells with an Neubauer-improved chamber (Paul Marienfeld GmbH & Co. KG, Lauda-Königshofen, Germany). The flocculation rate was determined using the Helm’s test according to Smart et al. (D’Hautcourt & Smart, 1999) with minor changes. Deviating from the original protocol, the test was performed in a volume of 4 mL and using a yeast cell number corresponding to OD_600_ = 10. During fermentation, pH was measured using a probe (Consort bvba, Turnhout, Belgium). For this, a beer sample was taken, and the yeast cells were removed from the wort medium by centrifugation at RT for 2 min at 4,000 rcf. For alcohol concentration and extract analysis, samples were centrifuged at RT for 2 min at 4,000 rcf and afterwards degassed in an ultrasonification bath (SONOREX DIGITEC DT 52 Ultrasonic bath, BANDELIN electronic GmbH & Co. KG, Berlin, Germany) for 15 min. The extract concentration of the wort medium was determined with a DMA5000 density meter (Anton Paar GmbH), while the alcohol concentration was measured using an Alcolyzer beer module (Anton Paar GmbH).

### 2.6 Analysis of the S. pastorianus flocculation proteome

#### 2.6.1 Yeast Protein Extraction

Yeast samples were taken 49 h, 72 h and 79 h after inoculation. For yeast with a non-flocculent phenotype, 40 mL of sample was taken while stirring. Flocculating yeast were let to rest unstirred for 15 min before 100 mL non-sedimented and 10 mL of sedimented yeast cells were sampled. Yeast cells were then collected by centrifugation for 2 min at 4 °C and 6,000 rcf and the cells were washed once with 20 mL ice cold sterile phosphate buffered saline (PBS) (Cat. No.: P4417-50TAB, Merck KGaA). Afterwards yeast cells were resuspended in 20 mL ice-cold PBS and transferred into a 30 mL plastic syringe (Cat. No: Z683647, Merck KGaA). Syringes were end-capped (Cat. No.: LBLL70, Becton Dickinson, Franklin Lakes, New Jersey, USA), placed into 50 mL conical centrifuge tubes and centrifuged at 4 °C and 6,000 rcf for 2 min. The supernatant was discarded and the yeast cells were then pressed into liquid nitrogen to snap-freeze and the cell pellets were stored at-80 °C until further processing.

Snap frozen yeast cells were grinded in liquid nitrogen using a mortar and pestle. Yeast powder was then transferred into a 5 mL reaction tube prefilled with 2 mL ice cold 20% (w/v) trichloroacetic acid (TCA, Cat. No. 10775151, Thermo Fisher Scientific) in 80% (v/v) acetone (Cat. No. 1070212511, Merck KGaA) and incubated at-20 °C overnight. Proteins were recovered by centrifugation at 6,000 rcf and - 10 °C for 20 min. The supernatant was removed and 2 mL ice cold 100 % acetone (Merck KGaA) was added, and samples were incubated at-20 °C for 30 min with intermittent vortexing every 10 min. Afterwards, the samples were centrifuged at 6,000 rcf,-10 °C for 20 min (Model DH.WCF00005, Witeg, Wertheim, Germany. The resulting pellet was resuspended in 2 mL lysis buffer (8 M urea (Cat. No. 1.08487.0500, Merck KGaA), 50 mM ammonium bicarbonate (AmBic, Cat. No. 09830-500G, Merck KGaA), pH 8.5) and proteins were solubilized using an ultrasonic rod (Model 250, Branson, Brookfield, CT, USA). For this, the sample were placed in an ice-cooled water bath and underwent two cycles of sonification with each cycle consisting of a 30 s pulse (10% amplitude) with a 30 s pause in between the two pulses to allow for sample re-cooling. Afterwards, cell debris and other insoluble materials were removed by centrifugation at 6,000 rcf and RT for 30 min. Afterwards, the supernatant was transferred into a fresh 2 mL tube and stored at-80 °C until further processing. Protein concentration was determined via a BCA Protein Assay Kit (Cat. No.: 23225, Thermo Fisher Scientific) according to manufacturer’s instructions.

#### 2.6.2 Protein digestion and TMT labelling

50 µg protein from each sample was reduced and alkylated for 45 min at RT using 10 mM Tris(2-carboxyethyl)phosphine hydrochloride (Cat. No.: C4706-2G, Merck KGaA) and 25 mM chloroacetamide (Cat. No.: C0267-100G, Merck KGaA). Afterwards, samples were diluted with 50 mM AmBic to 2 M urea and 1 µg trypsin protease (Cat. No.: 90058, Thermo Fisher Scientific) at a concentration of 1 µg/µL in 50 mM AmBic, pH 8.5 was added. Samples were incubated for 20 h at RT and digest was stopped by adding trifluoroacetic acid (TFA, Cat. No.: 1.08262.0025, Merck KGaA) to a final concentration of 1% (v/v). Peptides were desalted using in-house made C18 columns as described earlier (Jadav et al., 2023). After desalting, the peptides were freeze-dried (Alpha 2–4 LSC, Martin Christ Gefriertrocknungsanlagen GmbH, Osterode, Germany), resuspended in 71.4 mM HEPES pH 8.5 (Cat. No.: 54457-10G-F, Sigma-Aldrich, Saint Louis, Missouri, USA) (Zecha et al., 2019) and incubated in an ultrasonic bath for 15 min. Peptides were labelled with TMT10 plex label reagent set (Cat. No.: A44521, Thermo Fisher Scientific). The labelling reaction was stopped after 1 h by adding hydroxylamine (Cat. No.: 467804-10ML, Sigma-Aldrich) to a final concentration of 0.5% (w/v) and samples were incubated at RT for 15 min. Labelled peptides were then combined, snap frozen in liquid nitrogen and freeze dried. Peptides were resuspended in 2% (v/v) acetonitrile (Merck KGaA), 0.1% (v/v) formic acid (FA, Cat. No.: 5.33002.0050, Merck KGaA) and an additional desalting step using a C18 column was performed as above and samples were freeze dried. Finally, samples were resuspended in 12 µL 2% (v/v) acetonitrile (Merck KGaA), 0.1% (v/v) formic acid FA (Merck KGaA) and analysed via LC-MS/MS at the Vlaams Institute for Biotechnology (Gent, Belgium).

#### 2.6.3 LC-MS/MS analysis

Peptides (10 µl) were injected for LC-MS/MS analysis on an Ultimate 3000 RSLCnano system in-line connected to an Orbitrap Fusion Lumos mass spectrometer (Thermo). Trapping was performed on a 5 mm trapping column (300 μm internal diameter (I.D.), 5 μm beads, Cat. No. 11362113, Thermo Scientific) with a flow rate of 20 μl/min for 2 min in loading solvent A (0.1% (v/v) FA in water). Peptides separation was performed using a 110 cm µPAC™ Neo column Gen2 (Cat. No. 17897883, Thermo Fisher Scientific) kept at a constant temperature of 50 °C. Peptides were eluted using a two-component solvent system (MS solvent A (0.1% (v/v) FA in water) and MS solvent B (0.1% FA in acetonitrile)) with a linear gradient starting at 4% MS solvent B which reached 12% MS solvent B after 64 min. The gradient reached a MS solvent B concentration of 22% at 120 min, after 140 min 30% MS solvent B and at 90 min 70% MS solvent B. This was followed by a 10-minutes wash at 70% MS solvent B and column re-equilibration using 100% MS solvent A. The first 5 min of the elution gradient, the flow rate was set to 600 nl/min after which it was lower to 300 nl/min kept constant over the duration of the remaining gradient.

The mass spectrometer was operated in data-dependent positive ionization mode. Full-scan MS spectra (350-1,500 m/z) were acquired at a resolution of 120,000 in the Orbitrap analyzer with AGC target value of 4 x 10^5^, a maximum injection time of 50 ms and a cycle time of 3 seconds. The precursor ions (filtered for charge states 2-7, dynamic exclusion (60 s; +/- 10 ppm window) and minimal intensity of 5 x 10^4^) were selected in the quadrupole with an isolation window of 1.2 Da, accumulated to a target AGC of 1 x 10^4^ with a maximum injection time of 120 ms and fragmented using HCD with a 40% NCE. The product ions were analyzed in the orbitrap at a resolution of 50,000 (first mass 100 m/z). For internal calibration (lock mass) the polydimethylcyclosiloxane background ion at 445.120028 Da was used while instrument longitudinal performance was monitored via QCloud (Chiva et al., 2018; Olivella et al., 2021).

#### 2.6.4 Analysis of LC-MS/MS data

Raw data was searched using MaxQuant (version 1.6.3.4) (Tyanova et al., 2016) against a *Saccharomyces pastorianus* RefSeq database, downloaded on 25^th^ November 2020 from NCBI (10,760 entries) using standard settings. As fixed peptide modifications carbamidomethylation of cysteine was set, variable peptide modifications were set as oxidation of methionine and acetylation of the protein N-terminus. Trypsin was selected as protease, under omission of the proline rule. As run type “reporter ion MS2” was selected with isobaric labelling using TMT10plex. During first search the tolerance was set to 20 ppm and during main search a mass error tolerance of 4.5 ppm was used. Peptide and protein false discovery rates (FDR) were set to 5%. MaxQuant search results were further analysed using in-house written scripts (modified from (Khanam et al., 2021)) using R (version 4.5.1) (R Core Team, 2023). In brief, TMT reporter ion intensities of peptides which were quantified several times within a single LC-MS/MS analysis were averaged and resulting data was transformed and calibrated using variance stabilizing normalization (VSN) (Huber et al., 2002, 2003). Statistical analysis was conducted using limma (Ritchie et al., 2015) under application of robust hyperparameter estimation (Phipson et al., 2016). Peptides were regarded as differentially abundant if they exhibited an adjusted p-value ≤ 0.05 (5% FDR). Following R libraries were used: ggplot2 (Wickham, 2016), reshape2 (Wickham, 2007), vsn, limma, seqinr (Charif & Lobry, 2007), plyr (Wickham, 2011), stringr (Wickham, 2023), ggrepel (Slowikowski, 2024), ggpointdensity (Kremer, 2019), scales (Wickham et al., 2023), wesanderson (Ram & Wickham, 2023), matrixStats (Bengtsson, 2024) and viridisLite (Garnier et al., 2023). All mass spectrometry raw data and search engine results can be accessed with the identifier PXD081760 on ProteomeXchange (Deutsch et al., 2023) or JPST004805 on (Okuda et al., 2025). All R analysis scripts can be accessed via Zenodo (European Organization For Nuclear Research & OpenAIRE, 2013) under the DOI 10.5281/zenodo.21650315.

### 2.7 Construction of GFP drop-out vector

Yeast mutants overexpressing selected genes (*STE20* and *YPI1*) were constructed using the modular cloning MoClo Yeast Kit (YTK) (Cat. No.: Kit # 1000000061, Addgene, Watertown, Massachusetts, USA) (Lee et al., 2015).

First, *E. coli* containing the relevant plasmids (pYTK008, pYTK009, pYTK047, pYTK060, pYTK066, pYTK073, pYTK077, pYTK082, pYTK0083) were inoculated in 20 mL LB medium (Cat. No.: 113002065, MP Biomedicals, Santa Ana. California, USA) supplemented with the relevant antibiotic as per manufacturer’s manual. The culture was incubated overnight at 37 °C under stirring and afterwards centrifuged for 10 min at 4,000 rcf and RT. The supernatant was discarded and the cell pellet was resuspended in 200 µL buffer B1 (50 mM Tris (Cat. No.: 93350-1KG, Merck KGaA), 10 mM EDTA (Cat. No.: 324503-100GM, Merck KGaA), 100 μg/mL RNAse A (Cat. No.: EN0531, Thermo Fisher Scientific), pH 8) and vortexted. Afterwards, 200 μL B2 buffer (0.2 M sodium hydroxide (Cat. No.: 1064980500, Merck KGaA), 1% (w/v) sodium dodecyl sulphate (SDS, Cat. No.: 436143, Merck KGaA)) was added, mixed by inverting and incubated at RT for 5 min. This was followed by addition of 200 μL 5 M Potassium acetate (Cat. No.: 10522955, Thermo Fisher Scientific), pH 5.5) and the sample was mixed by inverting, which was followed by centrifugation at 4,000 rcf and RT for 15 min. The supernatant was transferred into a fresh 1.5 mL DNA LoBind tube (Cat. No.: 0030108051, Eppendorf, Hamburg, Germany) and 500 µL 100% ethanol (Cat. No.: 12498740, Thermo Fisher Scientific) was added. Sample was centrifuged at 21,000 rcf and-4 °C for 30 min and supernatant was discarded. 200 µL ultrapure water and 200 µL 8 M lithium chloride (Cat. No.: 310468, Merck KGaA) were added to the pellet and mixed by pipetting. The sample was then incubated at RT for 20 min, before being centrifuged at 21,000 rcf and 4 °C for 30 min. The supernatant was transferred into a fresh 1.5 mL DNA LoBind tube and mixed with 300 µL 100% isopropanol (Cat. No.: 10315720, Thermo Fisher Scientific) by inverting. Sample was incubated at-20 °C for one hour. After centrifuging the sample for 30 min at 21,000 rcf and-4 °C, the supernatant was discarded. The pellet was washed twice with 200 µL 70% (v/v) ethanol, by centrifuging at 21,000 g, RT for 5 min. The plasmid pellet was airdried before it was resuspended in 50 µL ultrapure water and incubated at 65 °C for 10 min before stored at-80 °C until further usage.

To create the GFP drop out vector, the relevant components (pYTK008, pYTK047, pYTK073, pYTK077, pYTK082, pYTK0083) were isolated as described above. Plasmids were mixed in an equimolar concentration (170 fmol each), 0.5 μL each of BsaI (Cat. No.: NEB R3733S, New England Biolabs) and T4 ligase (Cat. No.: M2622S, New England Biolabs) were added, along with 2.0 μL 10X T4 ligase buffer, and the solution was diluted to 20 μL final volume. The GoldenGate reaction (Engler et al., 2008) was performed overnight ([37 °C for 1 min, 16 °C for 1 min] x 30, 65 °C 5 min) in a Thermocycler. Ligation reactions (1-2 μL) were transformed into competent *E. coli* DH5α (Cat. No.: C2987H, New England Biolabs) according to manufacturer’s protocol and inoculated on LB^Amp^ agar plates at 37 °C. After 1-2 days green colonies were screened by colony PCR using primers covering a GoldenGate assembly junction (primers 43 & 44 in **Supplementary Table 1**, from pYTK083 AmpR to pYTK008 ConLS’). The positive plasmids were digested with BsmBI (Cat. No.: NEB R0739S, New England Biolabs) and run on an agarose gel to confirm expected insert sizes; then sequenced and validated by whole plasmid sequencing performed by Plasmidsaurus (Cologne, Germany) using Oxford Nanopore Technology with custom analysis and annotation (see Supplementary Data 1).

### 2.8 Gene amplification

*S. pastorianus* TUM 34/70 colonies were grown on a YPD-agar plate from cryoculture for approx. 5 days at RT. A single colony was isolated and resuspended in 100 µL lithium acetate buffer (200 mM lithium acetate (Cat. No.: L6883, Merck KGaA), 1% (w/v) SDS (Merck KGaA) and incubated at 70 °C for 15 min. 300 µL 96% (v/v) ethanol (Cat. No.: 10428671, Thermo Fisher Scientific) was added and mixed by inverting. Cell lysate was centrifuged at 15,000 g, RT for 3 min and supernatant was discarded. DNA was taken up in 100 µL TE-buffer (10 mM Tris (Merck KGaA), 1 mM EDTA (Merck KGaA), pH 8.0). Cell debris was removed using a benchtop minicentrifuge (Cat. No.: 16617645, Thermo Fisher Scientific) and DNA concentration was measured via a nanophotometer (NP80 touch, IMPLEN, München, Germany). DNA (250 ng starting amount) was amplified with Phusion Hot Start High Fidelity Polymerase (Cat. No.: 10441338, Thermo Fisher Scientific) according to user manual using 5x Phusion HF buffer (Cat. No.: F18L, Thermo Fisher Scientific) in a T100 Thermal Cycler (Bio-Rad Laboratories, Inc., Hercules, California, USA) for 30 cycles. The annealing temperature was set to 53 °C for all primer pairs. The used primers (19, 20, 23 and 24) are listed in **Supplementary Table 1**.

Amplified DNA was isolated with a QIAprep PCR Purification Kit (Cat. No.: 28106, Qiagen, Venlo, Netherlands) according to manufacturer instructions and stored at-80 °C until further usage.

### 2.9 Plasmid construction

The kinase overexpression vector was constructed from the previously constructed plasmids pYTK-GFP (GFP drop-out) and the inserts pYTK-009 (containing the pTDH3 promoter), pYTK-060 (containing the C-terminal 3xFLAG-6HIS tag), pYTK-066 (containing tTDH1 terminator) and the purified DNA-amplicon (*STE20* or *YPI1*) via GoldenGate assembly. A master mix was prepared with 100 pmol pYTK-GFP and 200 pmol of the respective inserts and amplicon. The assembly was done using a T7 ligase (Cat. No.: NEB M0318S, New England Biolabs, Ipswich, Massachusetts, USA) according to manufacturer’s protocol. The modular cloning master mix was used immediately for transformation of *E. coli* DH5α (New England Biolabs) according to manufacturer’s instructions. Resulting colonies which did not appear green were picked and grown overnight in LB^Amp^ medium (200 µg/mL ampicillin) at 37 °C. Plasmids were isolated as described in *2.7 Construction of GFP drop-out vector* and validated by whole plasmid sequencing performed by Plasmidsaurus using Oxford Nanopore Technology with custom analysis and annotation (see **Supplementary Data 1**).

The pYTK plasmid with an insert encoding for three alanine (pYTK-3Ala control) was constructed by Vectorbuilder Inc, Chicago, IL, USA (https://en.vectorbuilder.com/vector/VB251019-1294nfk.html).

### 2.10 Determination of antibiotic concentration for growth inhibition of TUM 34/70

The required Geneticin disulphate G418 concentration for selection of transformed *S. pastorianus* was determined via a kill curve approach. One colony from a YPD agar plate was inoculated in 10 mL wort and incubated over night at 22 °C. Overnight culture was diluted 1:20 with 12 °P wort and 100 µL yeast suspension were plated on YPD agar plates containing different concentrations of Geneticin disulphate G418: 200 µg/mL, 150 µg/mL, 100 µg/mL, 75 µg/mL, 50 µg/mL, 30 µg/mL, 15 µg/mL, 10 µg/mL, 5 µg/mL and 0 µg/mL (see **Supplementary Figure 3**).

### 2.11 Yeast transformation

*S. pastorianus* TUM 34/70 was grown for five days on a YPD agar plate at RT, a single colony was picked and inoculated in 25 mL YPD contained in a 1 L glass bottle. The culture was stirred overnight at 200 rpm and 22 °C. The next morning a yeast suspension from the overnight culture was diluted to OD_600_ = 0.3 in a volume of 200 mL YPD and incubated at 30 °C while being stirred at 200 rpm. Yeast was grown until it reached an OD_600_ = 1.4, 100 mL were taken and yeast cells were harvested by centrifugation at 1,000 rcf, 4 °C for 5 min (supernatant was discarded). Cell pellet was resuspended in 10 mL sterile ultrapure water and centrifuged at 1,000 rcf for 5 min at 4 °C. This washing step was repeated once and the cell pellet was resuspended in 25 mL pretreatment buffer (0.1 M lithium acetate (Merck KGaA), 10 mM dithiothreitol (Cat. No.: CL00.0481.0005, ChemLab NV, Zedelgem, Belgium), 10 mM Tris (Merck KGaA), 1 mM-EDTA (Merck KGaA), pH 8). Cells were incubated in the pretreatment buffer on a tube roller mixer SRT6 (Stuart Equipment, Stone, United Kingdom) for 30 min at RT. Cell pellet was washed twice with ice cold 1 M Sorbitol (Cat. No.: 10459760, Thermo Fisher Scientific), with centrifuging at 1,000 rcf, 4 °C for 5 min. Finally, cell pellet was resuspended in 500 µL 1 M Sorbitol. 100 µL yeast suspension was mixed with 1 µg plasmid and transferred into a 0.2 mm gap electroporation cuvette (Cat. No.: 15542423, Thermo Fisher Scientific). After a 5 min incubation on ice, electroporation took place using the “Sh5” setting of a MicroPulser Electroporator (Bio-Rad Laboratories, Inc., Hercules, California, USA). After electroporation the yeast suspension was directly transferred into a 1:1 mix of 500 µL YPD and 500 µL 1 M Sorbitol for recovery at 22 °C, 300 rpm and 6 h in a ThermoMixer C (Eppendorf). Yeast cells were then collected via centrifugation for 3 min, at 12,000 rcf and RT. The supernatant was discarded and cells were resuspended in 100 µL YPD and spread on a YPD^G418^ (10 μg/mL Geneticin disulphate G418) agar plate with a sterile Drigalski spatula for selection.

### 2.12 Phenotypic validation

To determine the kinase/phosphatase effect on the flocculation phenotype five mutants (derived from individual colonies) containing the plasmids pYTK-STE20, pYTK-YPI1, pYTK-3Ala respectively as well as five *S. pastorianus* wild-type (WT) colonies were propagated and inoculated for beer fermentation.

One liter of sterile 12 °P wort was inoculated 15 x 10^6^ cells/mL *S. pastorianus* TUM 34/70 WT (n = 5). For the overexpression mutants (pYTK-STE20, pYTK-YPI1 and the pYTK-3Ala control plasmid (n = 5 each) the wort was additionally supplemented with 10 µg/mL Geneticin disulphate G418. The fermentation was carried out at 12 °C (while stirring at 200 rpm) for 96 h, with sampling points every 24 h to determine cell density, pH, extract and flocculation rate. Yeast sampling for phospho-proteome analysis was conducted 42 h after starting fermentation. At the end of the fermentation 20 µL yeast culture from the beer fermentation was imaged via microscopy (Model BX41, Olympus, Tokyo, Japan) to determine morphology.

### 2.13 Phospho-proteomic analysis of STE20 and YPI1 overexpression mutants

#### 2.13.1 Protein extraction, digestion, TMT labelling and high pH fractionation

Yeast was snap frozen as described in chapter 2.6.1 and cell pellets were grinded using a freezer mill (Spex 6875, Cole-Parmer Instrument Company, LLC, Vernon Hills, Illinois, USA) using following settings: Pre-cooling for 5 min, milling with a rate of “15” for 3 cycles of 3 min with intermittent cooling for 2 min. Yeast powder was transferred into a 5 mL reaction tube prefilled with 2 mL ice cold 20% (w/v) TCA in 80% (v/v) Acetone and proteins were extracted as described for the proteome analysis in chapter 2.6.1. Proteins were resuspended in 2 mL phospho-lysis buffer (8 M urea (Merck KGaA), 50 mM AmBic (Merck KGaA), 1% (v/v) Phosphate Inhibitor Cocktail 2 (Cat. No.: P5726-5ML, Merck KGaA), 0.1% (v/v) Microcystin (Cat. No.: 33893-1ML-R, Merck KGaA), pH 8.5). Proteins were solubilized using an ultrasonic rod (Model 120 Sonic Dismembrator, Thermo Fisher Scientific) using following settings: 30 pulses of 1 second length at 50% amplitude with 1 second pause in between, while samples were being cooled in an ice-water bath. Afterwards, samples were centrifuged for 30 min at 7,000 rcf and RT, supernatant was retained and stored at-80 °C until further usage. The protein concentration of the samples was measured using a Pierce^TM^ BCA Protein Assay Kit (Thermo Fisher Scientific) according to manufacturer’s instructions.

Phospho-peptides were enriched, fractionated and TMT labelled as detailed in (Jadav et al., 2023) using 1 mg protein of each sample and trypsin instead of trypsin/LysC. In brief, phospho-peptides were enriched using TiO_2_ beads (TitanSphere Phos-TiO 10 µm, Cat. No. 5010-21315, GL Sciences, Japan) in a 1 mg protein to 4 mg beads ratio. The enrichment was repeated once, the phospho-peptides labelled using TMT16plex, pooled and freeze-dried. The pooled peptides were resuspended in 22 µL 10 mM ammonium formate pH 10 and fractionated using high-pH RP-HPLC (Vanquish Flex, Thermo Fisher Scientific) and 57 fractions were collected. These 57 fractions were concatenated into 19 fractions, freeze-dried and analysed via LC-MS/MS at the Vlaams Institute for Biotechnology (Gent, Belgium).

#### 2.13.2 - LC-MS/MS analysis

Peptides were resuspended in 0.1% (v/v) TFA in water and injected (200 ng and 800 ng in repeated run) onto a Vanquish™ Neo UHPLC system in-line connected to an Orbitrap Astral mass spectrometer (Thermo Fisher Scientific). Peptides were trapped on a 5 mm C18 column (Pepmap, 300 μm internal diameter (I.D.), 5 μm beads, Cat. No. 11362113, Thermo Fisher Scientific) using a trap-and-elute workflow in combined control mode (maximum flow: 40 µl/min, maximum pressure: 800 bar) and column was washed using buffer A (0.1% (v/v) TFA in water). Separation of peptides was carried out on an in-house packed column (75 µm x 100 mm, ReproSil Saphir 1.5 µm silica particles (Dr. Maisch HPLC GmbH, Ammerbuch-Entringen, Germany)) equipped with an electrospray tip via a P-2000 Laser Based Micropipette Puller (Sutter Instruments, Novato, CA, USA) and kept at a temperature of 45 °C. Peptides were eluted by a two buffer gradient system (buffer A and buffer B) at 250 nl/min: 0 % to 26% buffer B (0.1% FA in acetonitrile) from minute 0 to 17, increase to 44% buffer B at minute 20, increase to 56% buffer B at minute 21 and kept constant for 4 minutes (300 nl/min), followed by column equilibration in Pressure Control mode (separation column: fast equilibration, maximum pressure of 1500 bar, equilibration factor = 2; Trap column: fast wash and equilibration, wash factor = 100) with 100% buffer A.

The Orbitrap Astral was operated in data-dependent positive ionization mode with 0.5 s cycle time. Full-scan MS spectra (375-2000 m/z) were acquired at a resolution of 60,000 in the Orbitrap analyzer with a target AGC value of 5 x 10^6^ and a maximum injection time of 3 ms. The precursor ions were selected in the quadrupole (charge states 2-6, dynamic exclusion for 12 s within a +/- 5 ppm window and an intensity threshold of 5 x 10^3^) using an isolation window of 0.5 Da and an AGC target of 1 x 10^4^ with a maximum injection time of 20 ms. Precursors were fragmented using HCD with 38% NCE, first mass was set to 100 m/z. Internal calibration (lock mass) was performed on the polydimethylcyclosiloxane background ion (445.120025 Da). QCloud was used to monitor instrument longitudinal performance.

#### 2.13.3 Data analysis

Mass spectrometry raw data was analysed using FragPipe (version 24.0) (Kong et al., 2017; Teo et al., 2021) against a *S. pastorianus* FASTA (10,699 entries), downloaded on 18^th^ June 2026 from www.uniprot.org and amended by FragPipe with common contaminants and decoys. Standard “TMT18-Astral” workflow was chosen with following modifications: As variable modification, phosphorylation of serine, threonine and tyrosine was set, for validation MSBooster with DIA-NN (Demichev et al., 2020) for spectra & retention time prediction was chosen (Yang et al., 2023) and validated using Percolator (Käll et al., 2007), PTMprophet was used to localize PTMs (Shteynberg et al., 2019), false discovery rate (FDR) filtering at 5% threshold was done using Philosopher (da Veiga Leprevost et al., 2020) and TMT reporter ion intensities were quantified using IonQuant (Djomehri et al., 2020). Ion, peptide-to-spectrum match and peptide FDR were set to 5%, protein inference was disabled.

Search engine results were analysed using in-house written scripts in R (version 4.5.1). In brief, the dataset was filtered for TMT-labelled phospho-peptides, and unique peptides detected several times within a fraction were averaged. Data was then VSN transformed, calibrated and statistical testing was performed using limma. Peptides were regarded as differentially abundant if they exhibited an adjusted p-value ≤ 0.05 (5% FDR). All mass spectrometry raw data and search engine results are available under the identifier PXD081792 via ProteomeXchange and JPST004806 via jPOSTrepo. Analysis scripts are available from Zenodo under the DOI 10.5281/zenodo.21645491.

## 3 Results & Discussion

### 3.1 Flocculation is delayed in S. pastorianus TUM 34/70 by glutamine/glutamic acid supplementation

Flocculation delay by nitrogen sources was determined by supplementing glutamine & glutamic acid, ammonium sulphate, proline and urea after 24 h of beer fermentation (n = 3). At the 48 h fermentation time-point a significantly smaller flocculation rate could only be detected under glutamine & glutamic acid supplementation condition as in comparison to the non-supplemented control (Supplementation group: 52.8% (± 7.63 SD), Control: 70.0% (±6.69 SD), p = 0.045, **Figure 1A**, indicated by a *). This effect was not detected in the other conditions (urea and proline), the lower flocculation rate in the ammonium sulphate condition might be better explained by being a side-effect as a strong pH drop was detected in this condition (**Figure 1B**). It also is apparent that the ammonium sulphate supplementation caused an artefactual increase in measured extract (**Figure 1C**). The flocculation suppression effect of glutamine & glutamic acid was lessened after 72 h (p = 0.0945) and after 96 h non-detectable (p = 0.2734) (**Figure 1A**). Importantly, an influence of glutamine & glutamic acid spiking on the fermentation performance (as measured by pH, extract and cell density) was not detectable (see **Figure 1B-D**). Together, these results are in accordance with earlier research where flocculation induction in *S. pastorianus* by lack of glutamine and glutamic acid was shown (Ogata, 2012).

**Figure 1.**
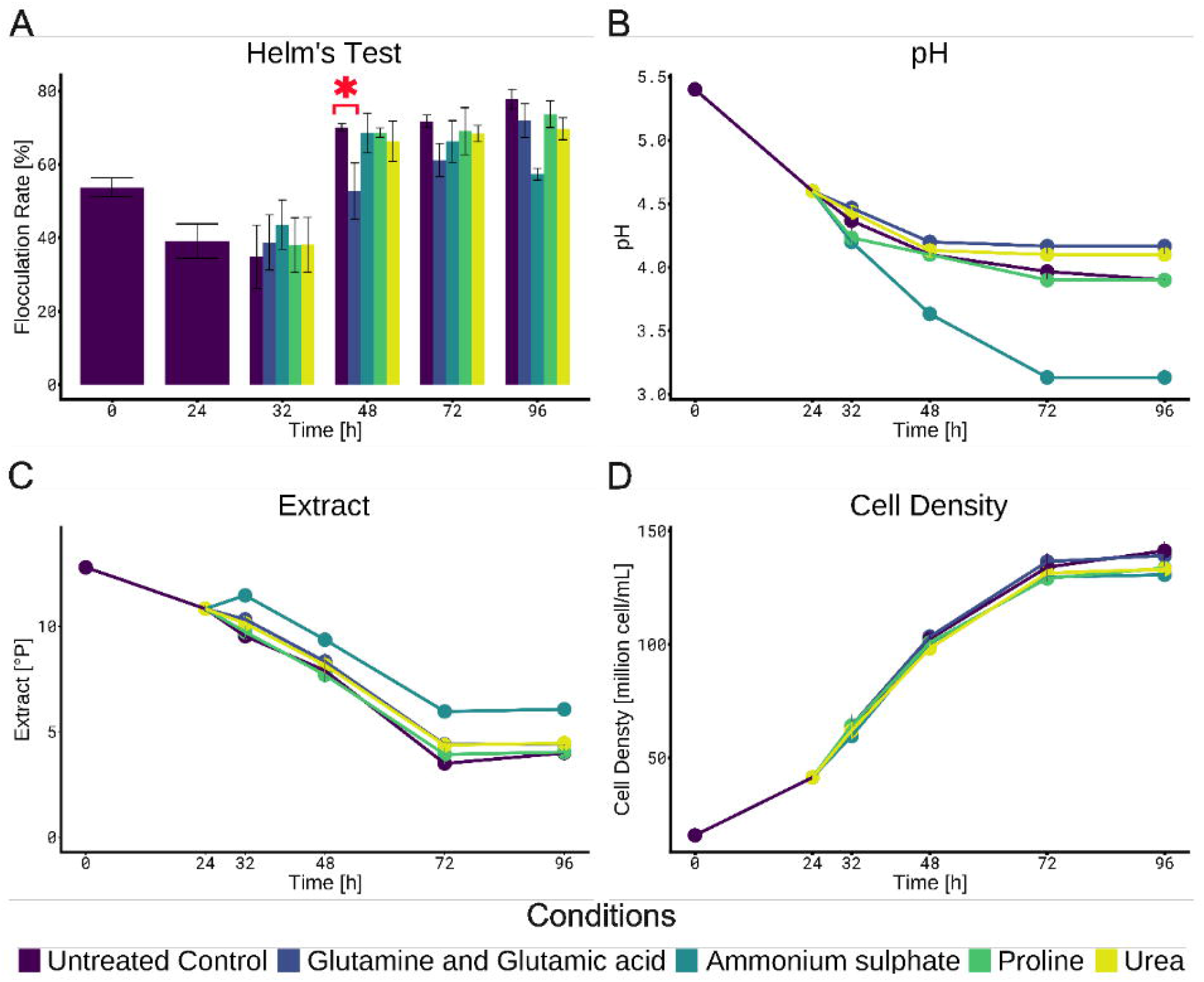
Influence of nitrogen sources on S. *pastorianus* TUM 34/70. (A) Results of Helm’s Test showing the different flocculation rates over the duration of a 96 h fermentation after supplementation with various nitrogen sources (glutamine & glutamic acid, ammonium sulphate, praline, urea and untreated control} at the 24 h timepoint. Red star indicates difference in flocculation rate between glutamine & glutamate supplementation as in comparison with untreated control (p = 0.045). (B) Development of pH, (C) extract (C) and (D) cell density after nitrogen supplementation. Error bars indicate standard deviations.

### 3.2 Proteome analysis of TUM 34/70 during flocculation

After having established that glutamine and glutamic acid supplementation delays flocculation in TUM 34/70, we were interested in which cellular events lead to the expression of this phenotype. For this, proteome analysis was conducted using yeast from glutamine & glutamate supplemented wort (n = 5) which was sampled 49 h (non-flocculent), 72 h (non-flocculent) and 79 h (flocculent) after start of fermentation, and a non-supplemented control (n = 5) which was sampled at 49 h (non-flocculent) and 72 h (flocculent). This setup allowed us to compare flocculent and non-flocculent yeast at several time-points to identify proteins which repeatedly showed differential abundance while exhibiting similar expression within the same phenotype and thus were potentially involved in flocculation (**Figure 2A**). As in the preliminary supplementation experiment, addition of glutamine and glutamic acid lead to temporary changes in flocculation rate as determined by the different cell densities in non-sedimented cells in supplemented and non-supplemented samples after 15 min of rest at the 72 h sampling point (p = 0.0454), with no visible influence on cell growth or extract reduction (see **Figure 2B and 2C**).

**Figure 2.**
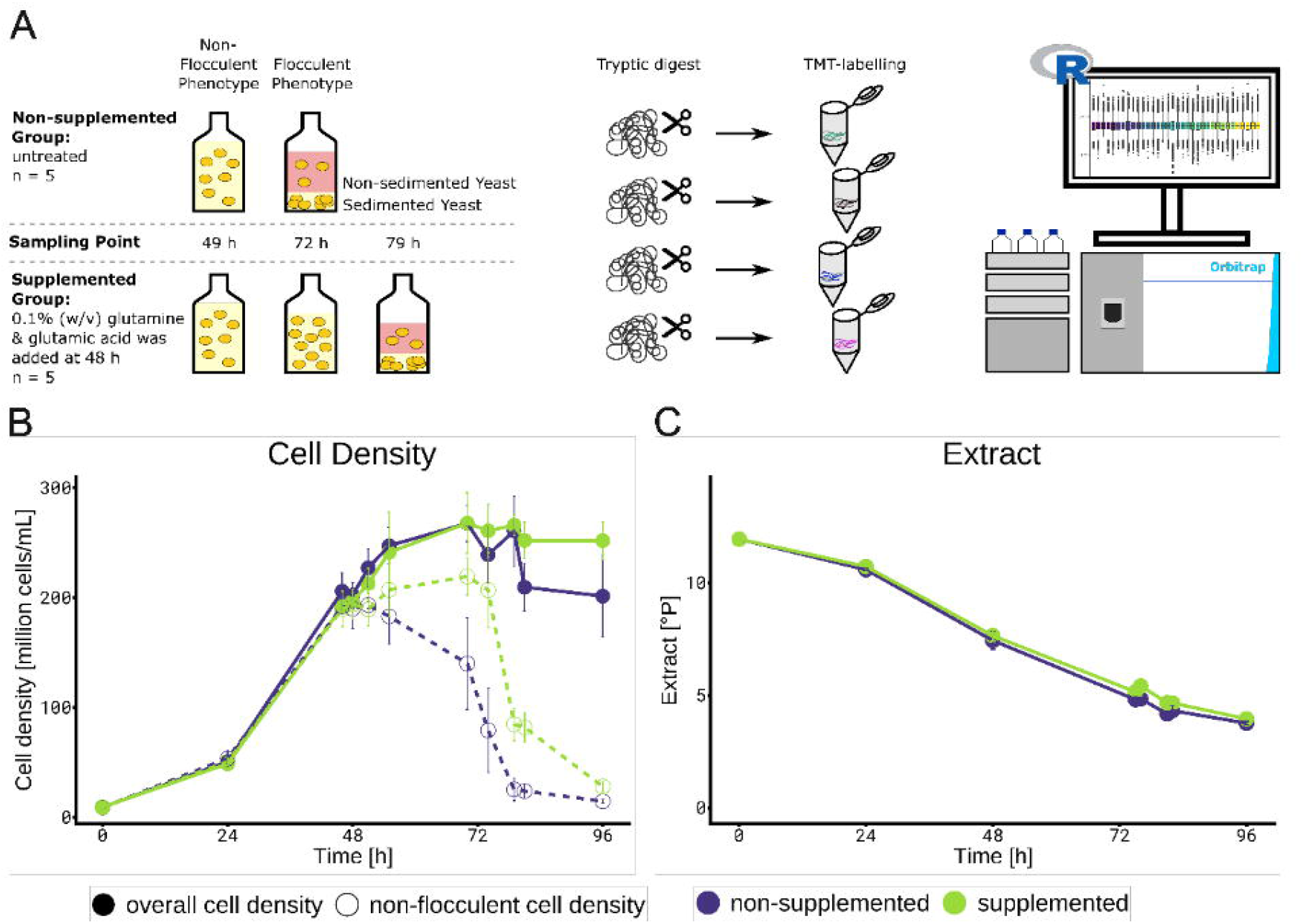
Proteome Analysis offlocculent and non-flocculent S. pastorianus TUM 34/70. (A) Experimental setup. Yeast (n = 5) was grown and supplemented after 48 h with 0.1% (w/v) glutamine and glutamic acid each and different cell populations were sampled after 49 h, 72 h and 79 h. Samples were processed for proteome analysis, identified and quantified via LC-MS/MS. (B) Growth curves indicating a lower flocculation rate (as per cell density of non-flocculent cells) in the supplemented group as in comparison with the non-supplemented control (p = 0.0454}. (C) Development of extract in growth medium during fermentation. Error bars indicate standard deviations.

In total, proteome analysis resulted in the identification and quantification of 101,903 peptides (24,445 unique) from 3,352 proteins (5% FDR). We laid focus on differentially abundant protein kinases and phosphatases as they are major signal-transduction enzymes (reviewed in (Ardito et al., 2017; Cohen, 1992)) and sustained phenotypes, such as flocculation, rely on a persistent cellular response which influences signal-transduction protein degradation profiles (reviewed in (Hunter, 2007; Lu & Hunter, 2009)). Based on the statistical analysis, the kinase *STE20_1* (QID79844, “Signal transducing kinases of the PAK”) which is a major kinase in the pseudohyphal growth pathway (Joshua et al., 2023; Liu et al., 1993) and further the *YPI1_2* phosphatase regulatory subunit (QID84787) were detected as higher abundant in flocculating cells (see **Table 1**). Although in the statistical testing of the supplemented vs. non-supplemented samples some of the adj. p-values of these proteins were higher than the usual 0.05 threshold, we still included them as they exhibited a small fold-change and higher adj. p-values might therefore be caused by a lack of statistical power. As next step, these two proteins were selected to validate their involvement in flocculation in *S. pastorianus* (complete proteomics results are shown in **Supplementary Table 2**).

**Table 1.** Differentially abundant peptides of Ste20p and Ypi1p during flocculation. QID gives the GenBank accession number. Adjusted p-value states the Benjamini-Hochberg corrected FDR. Positive logFC indicates higher abundant peptides in the first condition shown under Contrast (comparison the peptide was identified as differentially abundant). Non-Floc: Yeast showing overall non-flocculent phenotype, Sed.: Sedimented subpopulation of yeast after 15 min rest in overall flocculent sample, Non-Sed.: Non-sedimented subpopulation of yeast after 15 min rest in overall flocculent sample. S: Samples supplemented with glutamine & glutamic acid, NS: Non-supplemented samples. Numbers indicate proteomics sampling time point after inoculation.

| QID | Gene | Protein Description | Peptide Sequence | logFC | p-value | adj. p-value | Contrast |
| --- | --- | --- | --- | --- | --- | --- | --- |
| QID79844.1 | *STE20_1* | Signal transducing kinase of the PAK | GTNAAHEAGGYK | 1.19 | 0.002546 | 0.04248 | Sed.72.NS - Non.Floc.48.NS |
| QID79844.1 | *STE20_1* | Signal transducing kinase of the PAK | GTNAAHEAGGYK | 1.38 | 0.000839 | 0.03765 | Sed.72.NS - Non.Floc.48.S |
| QID79844.1 | *STE20_1* | Signal transducing kinase of the PAK | GTNAAHEAGGYK | 1.33 | 0.001416 | 0.06026 | Sed.72.NS - Non-Sed.72.NS |
| QID79844.1 | *STE20_1* | Signal transducing kinase of the PAK | GTNAAHEAGGYK | 0.92 | 0.012486 | 0.13450 | Sed.72.NS - Non-Floc.72.S |
| QID79844.1 | *STE20_1* | Signal transducing kinase of the PAK | GTNAAHEAGGYK | 0.79 | 0.028093 | 0.19374 | Sed.72.NS - Non-Sed.79.S |
| QID84787.1 | *YPI1_2* | Type1 phosphatases regulator ypi1 | NQTNM(ox)GSEQQQTAGSR | 2.65 | 0.000827 | 0.02477 | Sed.72.NS - Non.Floc.48.NS |
| QID84787.1 | *YPI1_2* | Type1 phosphatases regulator ypi1 | NQTNMGSEQQQTAGSR | 2.52 | 0.000082 | 0.00997 | Sed.72.NS - Non.Floc.48.NS |
| QID84787.1 | *YPI1_2* | Type1 phosphatases regulator ypi1 | NQTNM(ox)GSEQQQTAGSR | 3.33 | 0.000045 | 0.01054 | Sed.72.NS - Non.Floc.48.S |
| QID84787.1 | *YPI1_2* | Type1 phosphatases regulator ypi1 | NQTNMGSEQQQTAGSR | 2.44 | 0.000080 | 0.01460 | Sed.72.NS - Non.Floc.48.S |
| QID84787.1 | *YPI1_2* | Type1 phosphatases regulator ypi1 | NQTNM(ox)GSEQQQTAGSR | 3.23 | 0.000158 | 0.02649 | Sed.72.NS - Non-Sed.72.NS |
| QID84787.1 | *YPI1_2* | Type1 phosphatases regulator ypi1 | NQTNMGSEQQQTAGSR | 2.73 | 0.000044 | 0.01565 | Sed.72.NS - Non-Sed.72.NS |
| QID84787.1 | *YPI1_2* | Type1 phosphatases regulator ypi1 | NQTNMGSEQQQTAGSR | 1.82 | 0.001878 | 0.06095 | Sed.72.NS - Non-Sed.79.S |
| QID84787.1 | *YPI1_2* | Type1 phosphatases regulator ypi1 | NQTNMGSEQQQTAGSR | 1.63 | 0.004774 | 0.08644 | Sed.72.NS - Non-Floc.72.S |
| QID84787.1 | *YPI1_2* | Type1 phosphatases regulator ypi1 | ATQEPTRTQEAR | 1.24 | 0.039265 | 0.17405 | Sed.72.NS - Non-Floc.48.NS |
| QID84787.1 | *YPI1_2* | Type1 phosphatases regulator ypi1 | ATQEPTRTQEAR | 1.65 | 0.012274 | 0.14452 | Sed.72.NS - Non-Sed.72.NS |
| QID84787.1 | *YPI1_2* | Type1 phosphatases regulator ypi1 | NQTNMGSEQQQTAGSR | 1.83 | 0.010900 | 0.12891 | Sed.72.NS - Non-Sed.79.S |

### 3.3 Ste20p and Ypi1p are involved in establishing flocculation in S. pastorianus

As first approach, CRISPR/Cas-9 mediated knock-out (Gorter de Vries et al., 2017) of the *STE20* gene was attempted. Knock-out of *YPI1* was not attempted as this gene was described as essential in previous experiments in *S. cerevisiae* (García-Gimeno et al., 2003). However, the knock-out strategy was not successful (data not shown), and we decided to use a gene overexpression approach for *STE20* and *YPI1* instead.

Plasmids were introduced in *S. pastorianus* TUM 34/70 and successful transformation was validated by resistance to G418 and plasmid sequencing (**Supplementary Data 1**). Five colonies of yeast containing either *STE20*, *YPI1* or a control plasmid encoding for three alanine (3 Ala) instead, as well as five WT yeasts were propagated and used in beer fermentation for 96 h at 12 °C. Flocculation rate was determined every 24 h as function of remaining extract to control for natural variance in fermentation progression speed and a regression line was fit to determine systematic differences in flocculation rates between the experimental groups. It is important here to mention that the applied Helm’s test involves an EDTA wash step to be able to measure calcium-dependent flocculation, which also removes sugars attached to the Lg-Flo1p lectin. As such the actual flocculation rate of the yeast sample can be measured without masking from simple sugars in the growth medium which suppress the visual establishment of this phenotype in beer samples during fermentation (Dengis et al., 1995; Stratford & Assinder, 1991).

A sustained increase in flocculation rate as inverse function of extract remaining in the growth medium during fermentation between the 3 Ala control and *STE20* mutants (p = 6 x 10^-6^) and the *YPI1* mutants (p = 9 x 10^-7^) were detected, while no difference between the 3 Ala control and WT could be established (p = 0.349, **Figure 3A**). Importantly, no defects in extract reduction over time were observed in both overexpression mutants (**Figure 3B** and **3C**), although they showed a small lag in cell growth versus WT at the 72 h timepoint (**Figure 3D** and **3E**). Together, these results confirm the functional involvement of Ste20p and Ypi1p in flocculation and we were interested in the cellular pathways these two proteins act in. For this, we conducted a phospho-proteomic screen to establish their protein targets.

**Figure 3.**
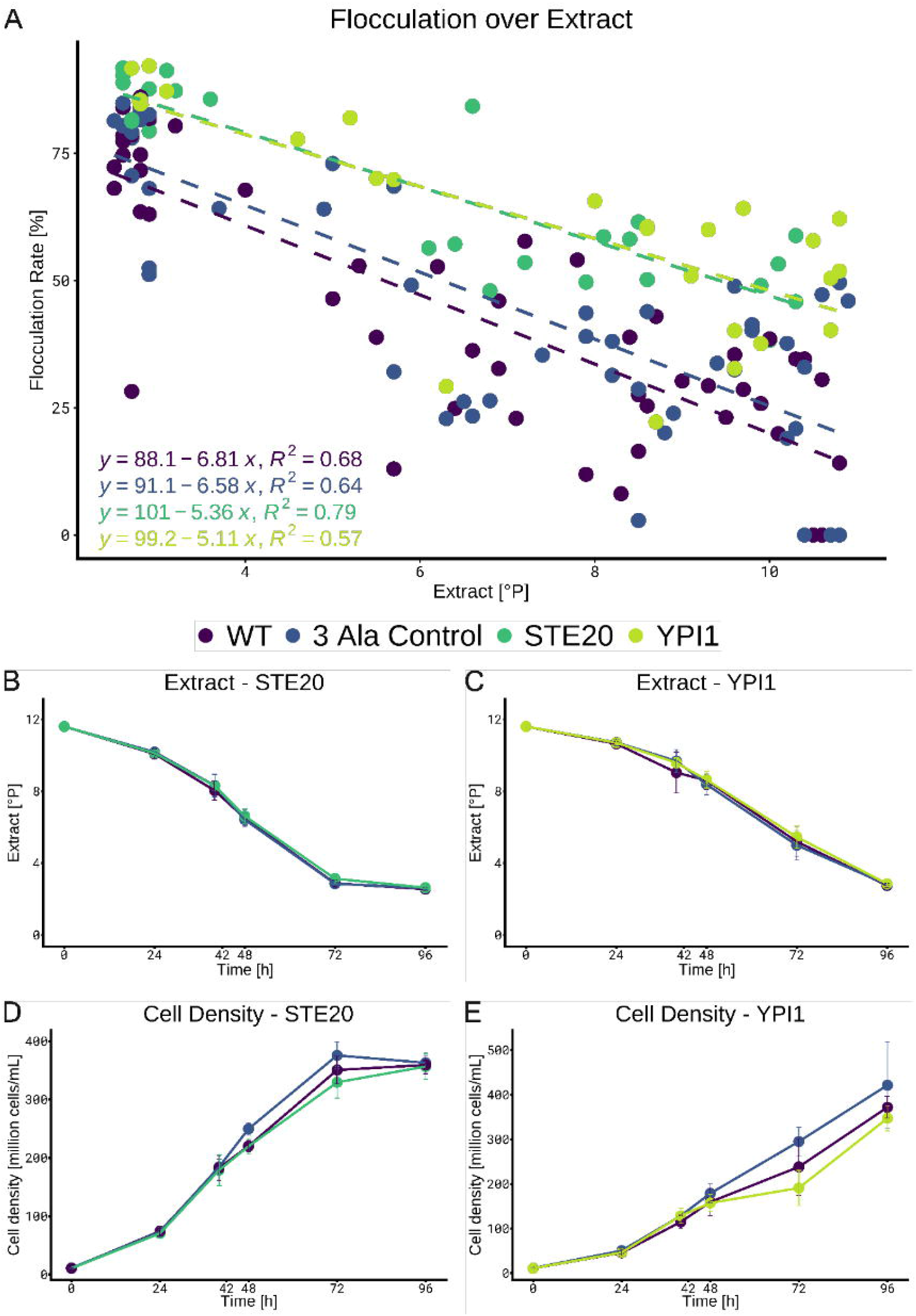
*STE20* and *YPl1* overexpression increase flocculation rate. (A) Linear regression of flocculation rate as a function of extract contained in growth medium between 24 hand 96 h after inoculation in S. *pastorianus* TUM 34/70 *WT,* 3 Ala control, *STE20* and *YPl1* overexpression. Development of extract contained in growth medium for (B) *STE20* overexpression and (C) *YP/1* overexpression vs *WT* and 3 Ala control (n = 5 for all). (D) and (E) show cell growth in *STE20* and *YP/1* overexpression, respectively, in comparison with WT and 3 Ala control (n = 5 for all). Error bars indicate standard deviations.

### 3.4 Ste20p and Ypi1p repurpose the pseudohyphal growth pathway for flocculation

For the phosphoproteomic screening, yeast cells were harvested at the 42 h fermentation timepoint (Figure 3B-E) and for the *STE20* overexpression experiment 33,280 unique phospho-peptides (5% FDR) from 4,766 proteins were identified, while for *YPI1* 34,285 unique phospho-peptides (5% FDR) from 5,288 proteins were detected (in total 51,413 unique phospho-peptides (5% FDR) corresponding to 36,896 phosphorylation sites (2,268 sites ≤ 1% FLR) from 6,372 proteins). Comparison of 3 Ala control vs WT (n = 5 for both conditions) resulted in a single differentially abundant phospho-peptide in case of *STE20-* and no differences for *YPI1* overexpression (n = 5 for both conditions, 5% FDR). As such, also due to the non-detectable difference in flocculation rate between WT and 3Ala control, these groups were regarded as equivalent, and as such combined in the statistical analysis. In case of *STE20* overexpression, 218 higher abundant phospho-peptides were detected (5% FDR) belonging to 22 proteins (**Table 2**). The majority of the identified peptides belonged to Ste20p, giving evidence for successful overexpression of this protein (**Figure 4A**). As Ste20p is a kinase involved in pseudohyphal growth, we next checked if any of the higher abundant phospho-peptides belonged to proteins also involved in this biological process. Out of the 22 detected proteins the orthologues of 4 proteins (exclusive Ste20p) were earlier reported to be also involved in pseudohyphal growth: Sok2p (Pan & Heitman, 2000; Ward et al., 1995), Vip1p (Norman et al., 2018; Pöhlmann & Fleig, 2010), Reg1p (Orlova et al., 2006) and Ptp1p (Fasolo et al., 2011) (**Figure 4A and Table 2**). Filamentous growth in *S. cerevisiae* is also controlled via peptide pheromones (Liu et al., 1993) and a phospho-peptide of the pheromone-regulated membrane protein 5 (Prm5p) was also identified to be higher abundant under *STE20* overexpression conditions (**Figure 4A** and **Table 2**). Interestingly, 2 proteins (Pbs2p and Bni4p, see **Figure 4A** and **Table 2**) which had lower abundant phospho-peptides under *STE20* overexpression are also (indirectly) related to pseudohyphal growth. Pbs2p is targeted by Ste11p under hyperosmotic stress (Maeda et al., 1995) and Ste11p is involved in pseudohyphal growth (Kim & Rose, 2015; Liu et al., 1993; Lorenz & Heitman, 1997). On the other hand, the protein Bni4p is targeted by the phosphatase Glc7p (Walsh et al., 2002) which in turn is targeted by Ypi1p (García-Gimeno et al., 2003). Interestingly, a phospho-peptide of the gene *STE20*_2 was detected as lower abundant under *STE20*_1 overexpression (see **Table 2**), however the implications are currently unclear and would be interesting to study.

**Figure 4.**
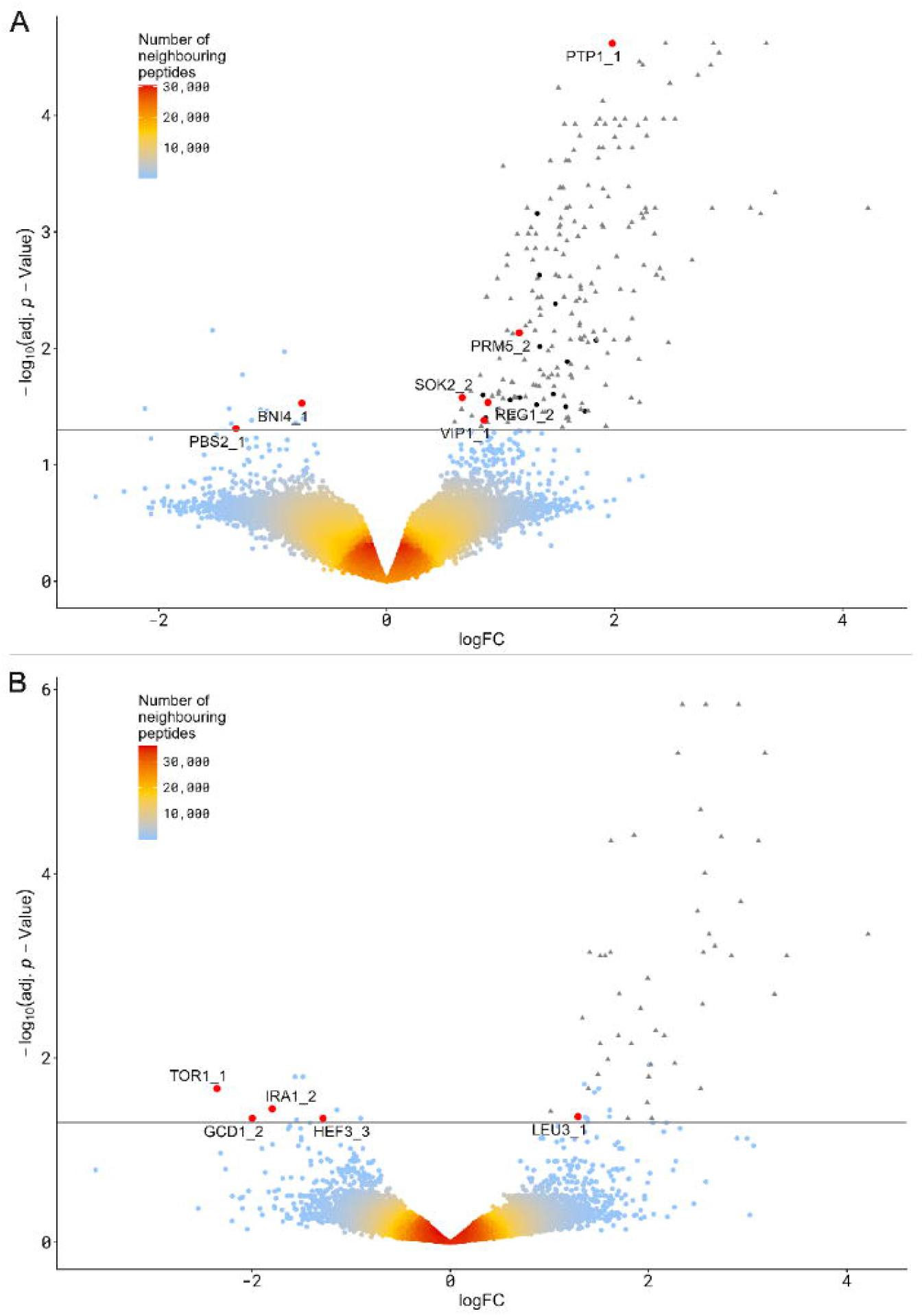
Phosphoproteomics of *STE20* and *YPl1* overexpression. Volcano plot analysis indicating phospho-peptides. Phospho-peptides with a positive logFC are higher abundant under (A) *STE20* overexpression (as in comparison with WT and 3 Ala control), grey triangles display identified Ste20p phospho-peptides indicating successful overexpression and (B) for *YPf1* overexpression, while grey triangles here indicate Ypi1p phospho-peptides. Gene names of proteins involved in the pseudohyphal growth pathway are labelled. Horizontal line indicates a 5% FDR.

**Table 2.**
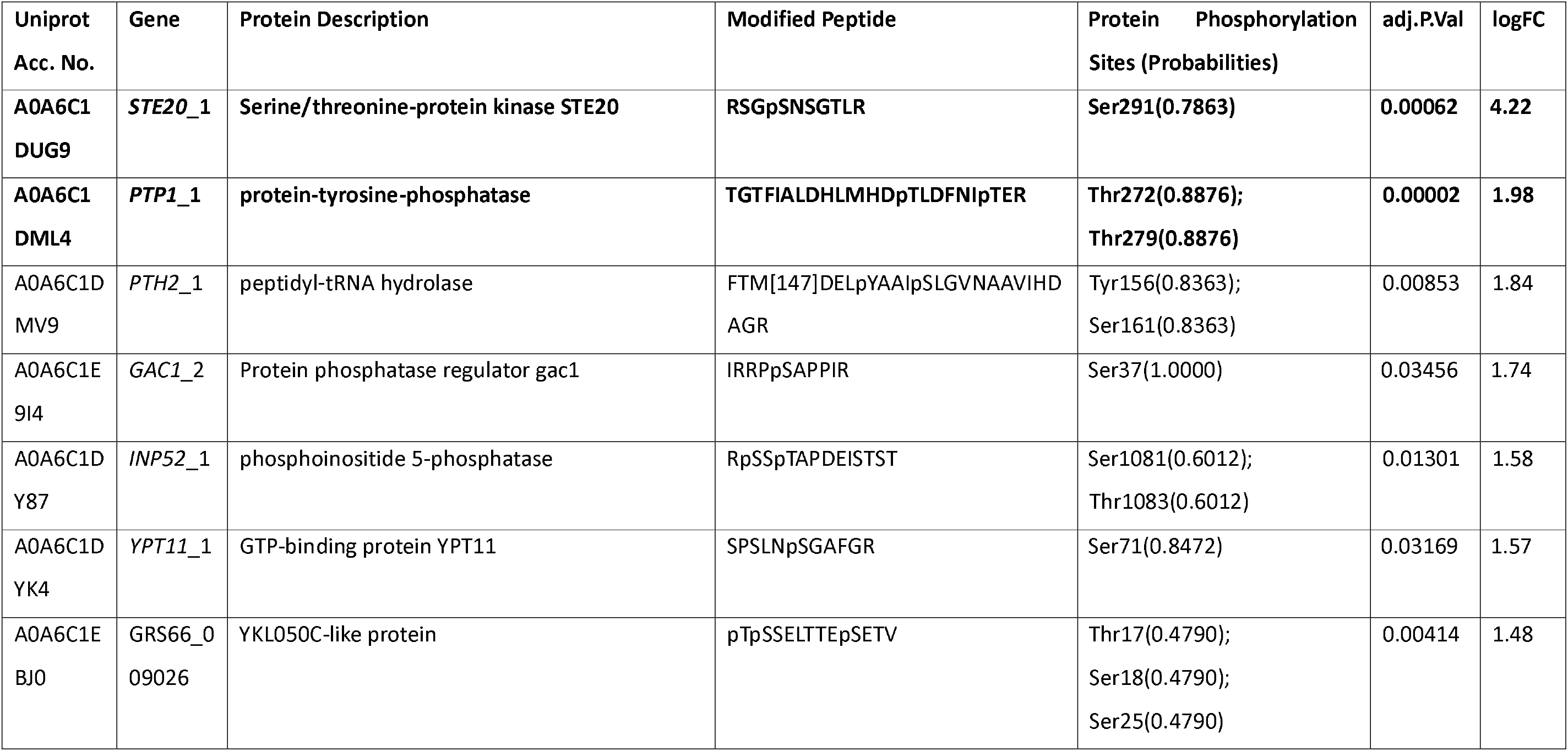
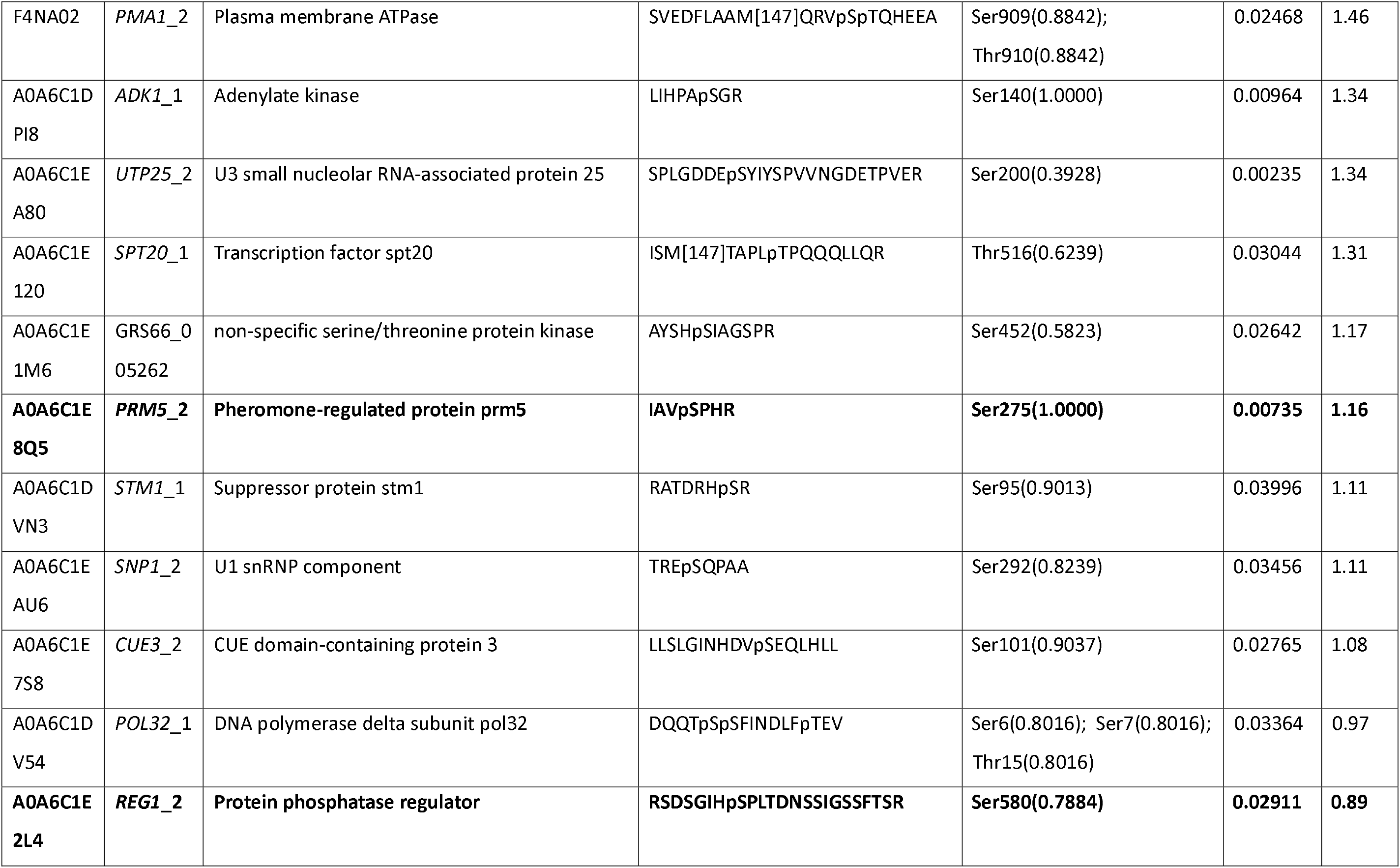
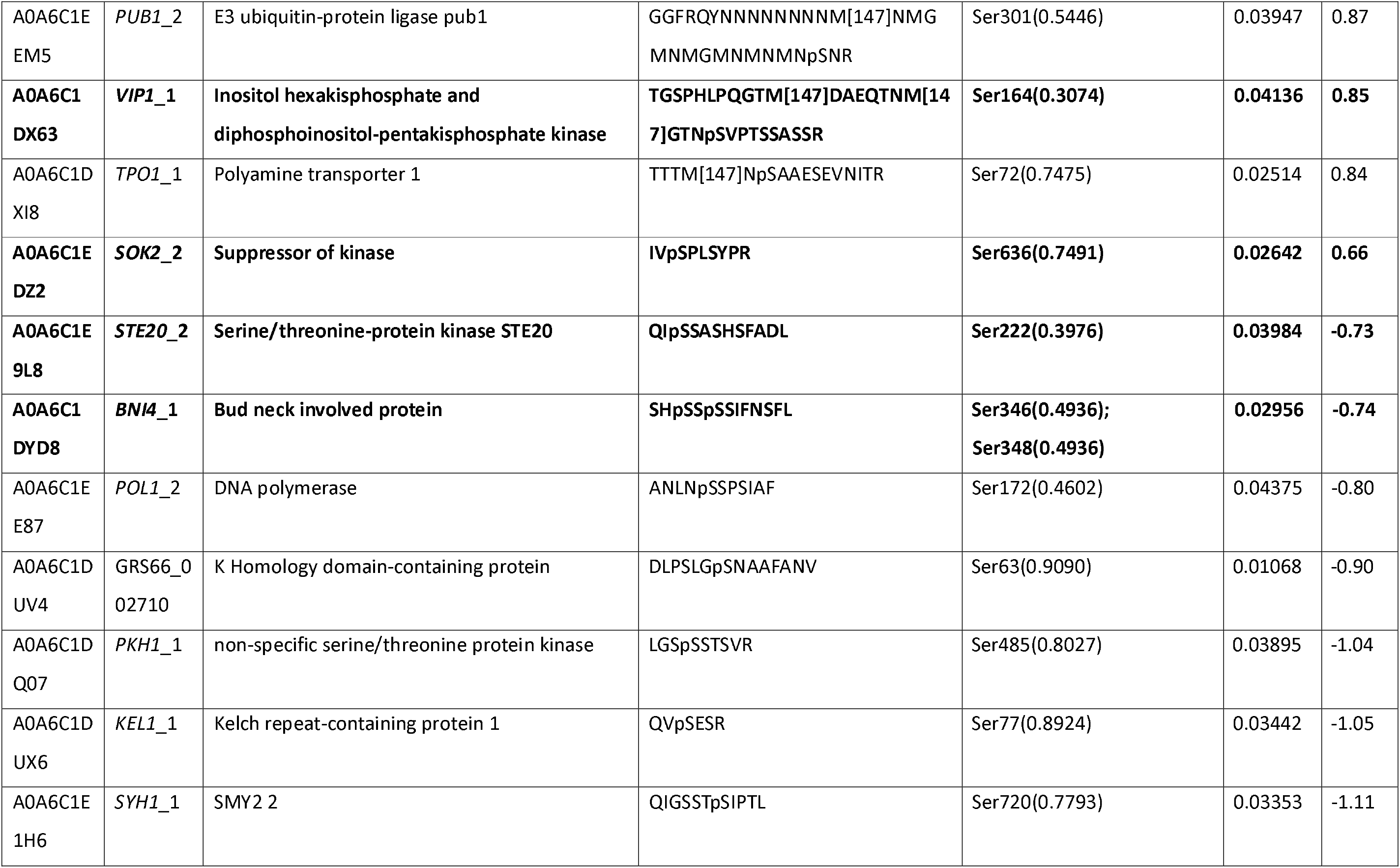
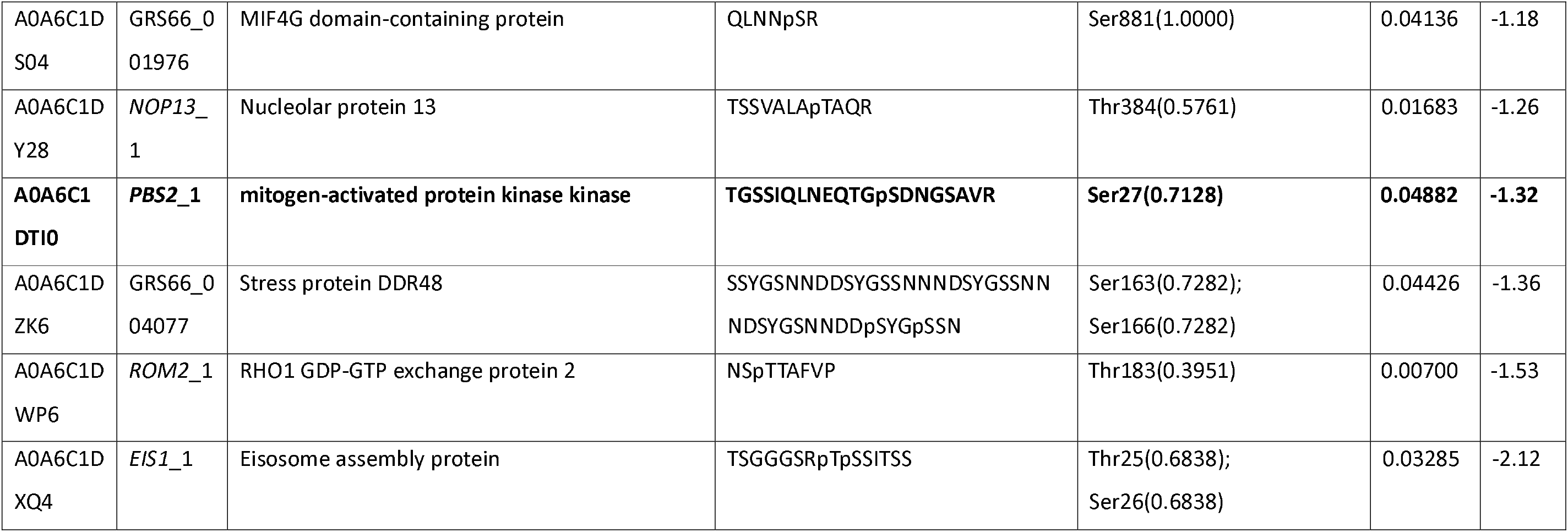
Differentially abundant phospho-peptides under STE20 overexpression. Uniprot Acc. No. gives the Uniprot accession number of the protein. “Peptide” column displays the identified amino acid sequence, M[147] indicates an oxidized methionine, the most probable phosphorylation site is indicated by a preceding “p”. Column “Protein Phosphorylation Sites (Probabilities)” indicates the localization of the phosphorylations site within the protein, probability of localization being correct within the identified peptide sequence is given in brackets. Adjusted p-value states the Benjamini-Hochberg corrected FDR. Positive logFC indicates a higher abundant peptide under STE20 overexpression. In case several phospho-peptides of the same protein were identified, only the one with the highest absolute fold change was retained (see also **Figure 4A**). Genes described to be involved in pseudohyphal growth/amino acid starvation response are given in bold.

Also, in case of *YPI1* overexpression the majority of higher abundant peptides belonged to Ypi1p which indicates successful overexpression of the gene (see **Figure 4B**). As Ypi1p is a phosphatase inhibitor for Glc7p (García-Gimeno et al., 2003), the phospho-peptides detected as higher abundant under *YPI1* overexpression were of primary interest. Only a single protein (Leu3p, Uniprot Acc. No: A0A6C1DXL2) connected to pseudohyphal growth (Shively et al., 2013) exhibited a higher abundant phospho-peptide. However, a total of 9 lower abundant phospho-peptides, belonging to 9 proteins, were detected (**Table 3**). Several orthologues of these proteins are reported to be either involved in pseudohyphal growth (Ira1p (Harashima et al., 2006), Hef3p (Laxman & Tu, 2011)) and/or amino acid starvation response (Tor1p (Wilson & Roach, 2002), Gcd1p (Hinnebusch & Fink, 1983) and Hef3p (Laxman & Tu, 2011)), which is a known trigger for pseudohyphal growth (Britton et al., 2025; Gimeno et al., 1992; Lengeler et al., 2000; Lodolo et al., 2008; Mösch, 2000; Mösch & Fink, 1997) (**Figure 4B** and **Table 3**). This higher occurrence of proteins involved in pseudohyphal growth and amino acid starvation response having phospho-peptides being lower abundant is a surprising result and hints that Ypi1p might have a complex influence on its downstream signaling phosphatase/kinase network.

**Table 3.**
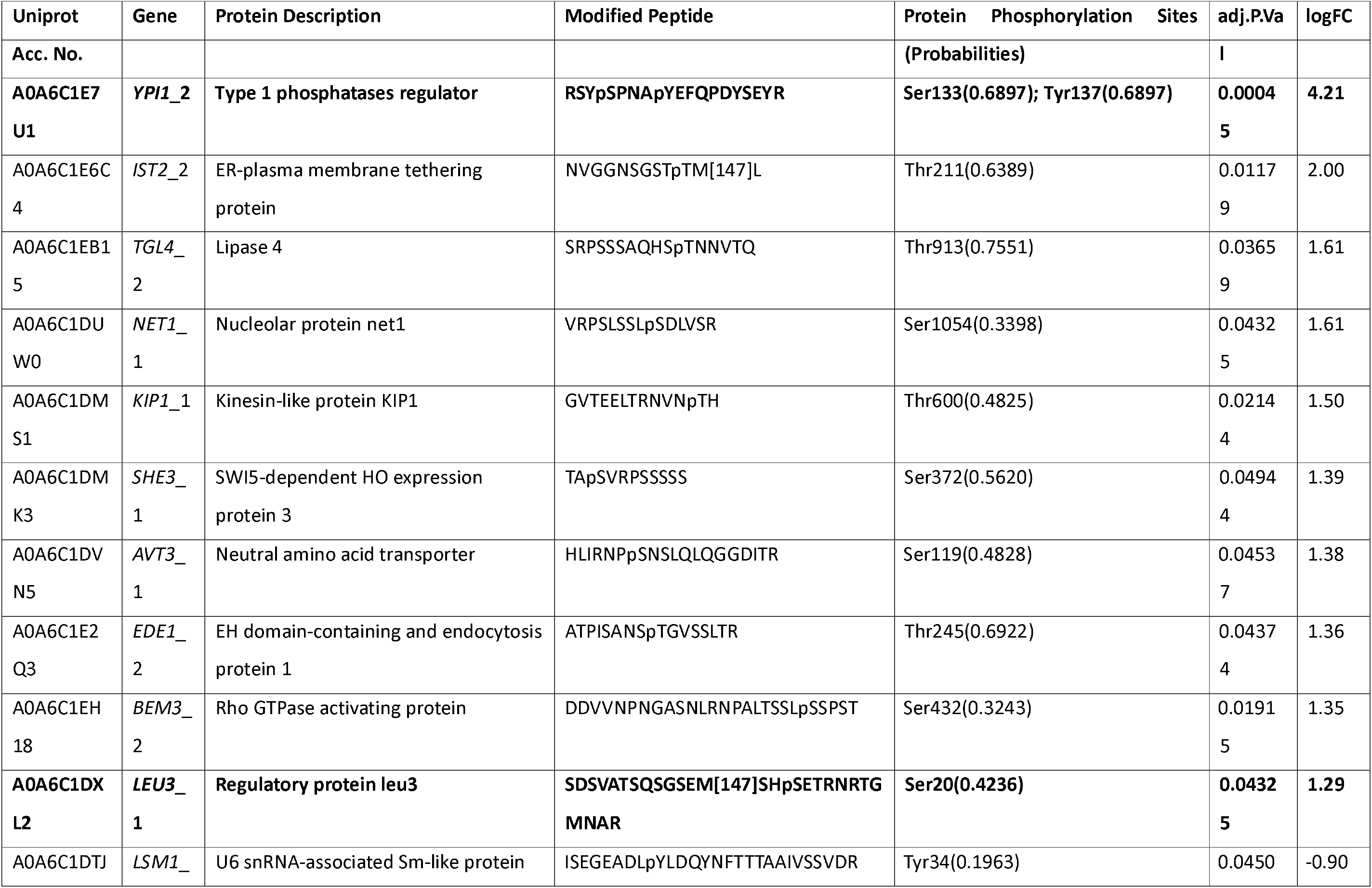
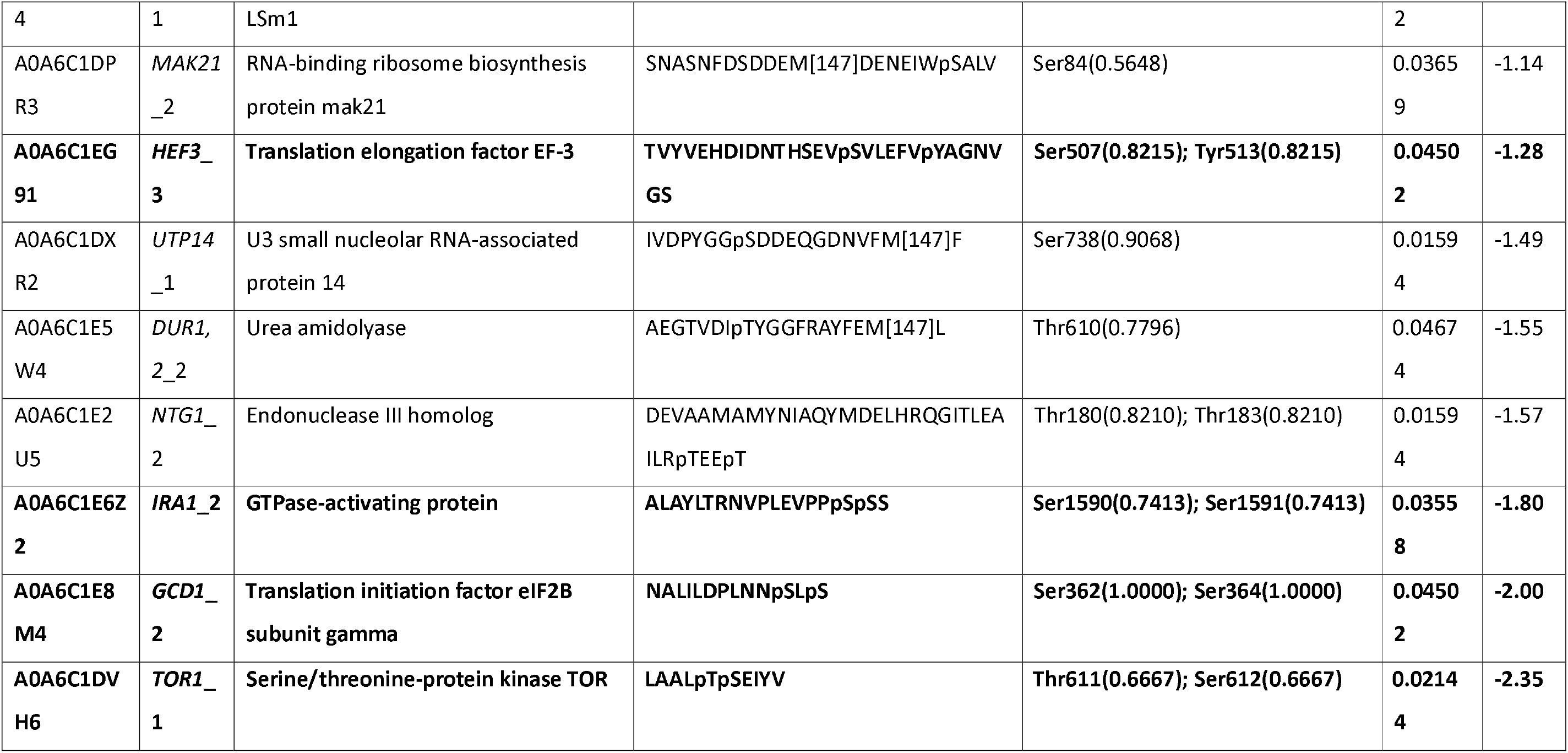
Differentially abundant phospho-peptides under YPI1 overexpression. Uniprot Acc. No. gives the Uniprot accession number of the protein. “Peptide” column displays the identified amino acid sequence, M[147] indicates an oxidized methionine, the most probable phosphorylation is site indicated by a preceding “p”. Column “Protein Phosphorylation Sites (Probabilities)” indicates the localization of the phosphorylations site within the protein, probability of localization being correct within the identified peptide sequence is given in brackets. Adjusted p-value states the Benjamini-Hochberg corrected FDR. Positive logFC indicates a higher abundant peptide under YPI1 overexpression. In case several phospho-peptides of the same protein were identified, only the one with the highest absolute fold change was retained (see also **Figure 4B**). Genes described to be involved in pseudohyphal growth/amino acid starvation response are given in bold.

Together, these data strongly suggest that Ste20p and Ypi1p are targeting proteins whose orthologues are involved in the pseudohyphal growth pathway. However, we did not detect differential phosphorylation of Flo11p (see **Supplementary Table 3**), which is a major protein in the establishment of pseudohyphal growth and for flocculation in *S. cerevisiae* var. *diastaticus* (Bayly et al., 2005; Chen & Fink, 2006; Lo & Dranginis, 1996; Ryan et al., 2012). Microscopy images of overexpression mutants did not show a pseudohyphal morphology (as defined by presenting filamentous pseudohyphae of elongated cells), but a flocculation morphotype after 72 h (**Figure 5A-D**), with a classical calcium-dependent flocculation type as per Helm’s test (**Figure 3B-E**). This is in unison with an earlier report that genes involved in pseudohyphal growth during nutrient limitation in *S. pastorianus* propagation are up-regulated, where also no such phenotype could be established (Gibson et al., 2010). Thus, our data are not consistent with pseudohyphal growth by Flo11p in case of *STE20* and *YPI1* overexpression, but compatible with the signaling network responsible for pseudohyphal growth in *S. cerevisiae* being repurposed in *S. pastorianus* (*S. cerevisiae* x *S. eubayanus*) to establish a flocculation phenotype instead.

**Figure 5.**
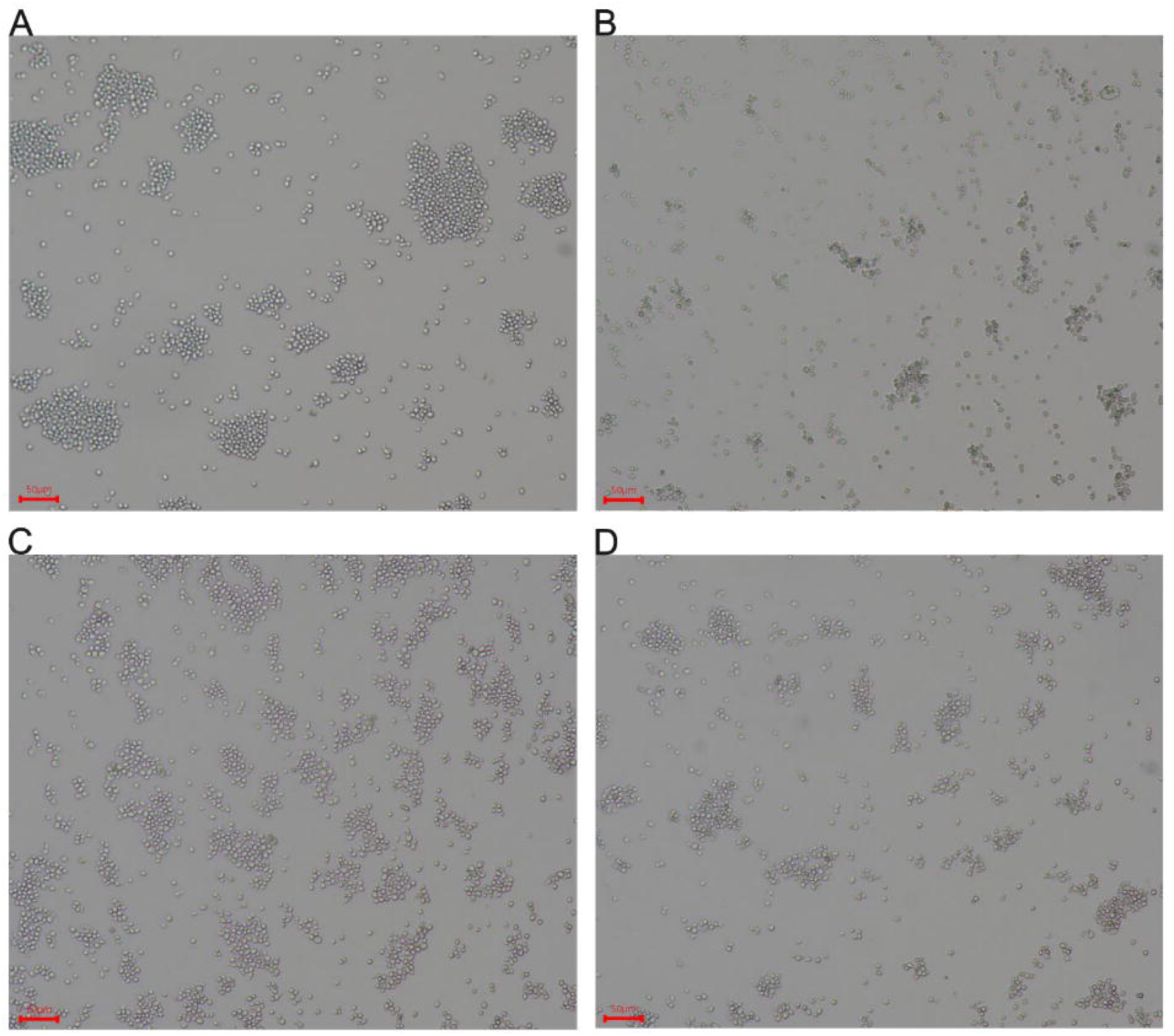
Microscopy images of S. pastorianusTUM 34/70. (A)TUM 34/70\NT, (B) 3 Ala control, (C) *STE20* overexpression and (D) *YP/1* overexpression taken 72 h after inoculation showing a flocculent phenotype with no indication of pseudohyphal growth. Bar indicates 50 µm length.

## 4 Conclusion

In summary, our preliminary experiments showed that flocculation in *S. pastorianus* can be delayed by supplementing a preferred nitrogen source (glutamine and glutamic acid), which is an extension of earlier results showing the induction of flocculation by amino acid starvation (Ogata, 2012). Further, we established Ste20p and Ypi1p to be higher abundant during flocculation and we validated their functional involvement using overexpression mutants. Biological function analysis of Ste20p and Ypi1p established their protein targets (as reported for other yeast species) to be associated with pseudohyphal growth, however an establishment of such a phenotype in *S. pastorianus* could not be detected. Together, these data constitute first-time evidence that cellular pathways which are normally involved in pseudohyphal growth are repurposed in *S. pastorianus* to lead instead to flocculation. As such, these results will be directly applicable to investigate involvement in flocculation defects typically encountered in bottom-fermenting brewing yeasts over re-pitching cycles in the brewing industry and examine if e.g. only genetic re-arrangements in Lg-*FLO1* lead to this phenomenon (Powell et al., 2003; Sato et al., 2001, 2002).

However, our study has several limitations. Currently, it is unknown how these involved proteins establish a flocculation phenotype, i.e. if the chromosomal translocation of the flocculation gene Lg-*FLO1* (Ogata et al., 2008) in *S. pastorianus* is a main driving factor or if the specific *FLO11* variant is responsible for flocculation in TUM 34/70. While phospho-peptides of other proteins involved in flocculation (Flo5p, Flo8p, Flo9p) were also identified, they did not appear to be differentially abundant (adj. p-value > 0.05, see **Supplementary Table 3**) and as such it is inconclusive if they are 1) affected by either Ste20p or Ypi1p or 2) have lost their function for flocculation in *S. pastorianus*. Further, TUM 34/70 is a bottom-fermenting yeast of the Frohberg subtype, and it remains to be investigated if our results also apply in Saaz subtypes as they are characterized by different genomic rearrangements (Gallone et al., 2018; Salazar et al., 2019; Wendland, 2014). The kinase Ste20p, the phosphatase Ypi1p and their individual protein targets would also need to be more thoroughly characterized, especially as in our sets of differentially abundant phospho-peptides potential downstream kinase and phosphatases were detected and as such made e.g. (de)phosphorylation motif analysis for Ste20p and Ypi1p not applicable. For such functional studies a knock-in approach to render Ste20p catalytically inactive would be advantageous, e.g. to explicitly implicate the kinase function in flocculation signaling and would also leave the examined genes under their original promotor preventing spurious off-target phosphorylations caused by an overexpression setup. However, for this, reliable genomic tools for *S. pastorianus* would need to be developed as current CRISPR/Cas9 approaches are not sufficiently efficient (Bennis et al., 2023).

## CRediT author statement

**Katharina Seibel:** Conceptualization, Methodology, Investigation, Formal analysis, Software, Visualization, Writing - Original Draft, Writing - Review & Editing. **Eoin Ó Cinnéide:** Methodology, Investigation, Formal analysis, Writing - Review & Editing. **Riley Schmalhaus:** Investigation, Formal analysis, Writing - Review & Editing. **Mirjam Haensel:** Conceptualization, Supervision, Writing - Review & Editing. **Florian Weiland:** Conceptualization, Methodology, Formal analysis, Software, Visualization, Supervision, Project administration, Writing - Original Draft, Writing - Review & Editing, Funding acquisition.

## Acknowledgments

We would like to thank the proteomics core of the Vlaams Institute for Biotechnology (VIB) for conducting the LC-MS/MS analyses. Further, we thank Quinten Deparis from the lab of Kevin Verstrepen (KU Leuven) for helpful discussions regarding transformation protocols for *S. pastorianus* and the lab of Prof. Pascual Sanz (Instituto de Biomedicina de Valencia, Spain) for providing us the original plasmid for *YPI1* overexpression.

## References

Ardito, F., Giuliani, M., Perrone, D., Troiano, G., & Muzio, L. L. (2017). The crucial role of protein phosphorylation in cell signaling and its use as targeted therapy (Review). International Journal of Molecular Medicine, 40(2), 271–280. 10.3892/ijmm.2017.3036

Bayly, J. C., Douglas, L. M., Pretorius, I. S., Bauer, F. F., & Dranginis, A. M. (2005). Characteristics of Flo11-dependent flocculation in Saccharomyces cerevisiae. FEMS Yeast Research, 5(12), 1151–1156. 10.1016/j.femsyr.2005.05.004

Bengtsson, H. (2024). matrixStats: Functions that Apply to Rows and Columns of Matrices (and to Vectors). https://CRAN.R-project.org/package=matrixStats

Bennis, N. X., Kostanjšek, M., van den Broek, M., & Daran, J.-M. G. (2023). Improving CRISPR-Cas9 mediated genome integration in interspecific hybrid yeasts. New Biotechnology, 76, 49–62. 10.1016/j.nbt.2023.04.001

Britton, S. J., Niemetz, J., Haensel, M., White, J. S., Maskell, D. L., & Weiland, F. (2025). Quorum sensing in Saccharomyces cerevisiae brewing strains: Effects of 2-phenylethanol on proteomic, lipidomic, and metabolomic profile. FEMS Yeast Research, 25, foaf036. 10.1093/femsyr/foaf036

Charif, D., & Lobry, J. R. (2007). SeqinR 1.0-2: A contributed package to the R project for statistical computing devoted to biological sequences retrieval and analysis. In U. Bastolla, M. Porto, H. E. Roman, & M. Vendruscolo (Eds.), Structural approaches to sequence evolution: Molecules, networks, populations (pp. 207–232). Springer Verlag.

Chen, H., & Fink, G. R. (2006). Feedback control of morphogenesis in fungi by aromatic alcohols. Genes & Development, 20(9), 1150–1161. 10.1101/gad.1411806

Chiva, C., Olivella, R., Borràs, E., Espadas, G., Pastor, O., Solé, A., & Sabidó, E. (2018). QCloud: A cloud-based quality control system for mass spectrometry-based proteomics laboratories. PloS One, 13(1), e0189209. 10.1371/journal.pone.0189209

Cohen, P. (1992). Signal integration at the level of protein kinases, protein phosphatases and their substrates. Trends in Biochemical Sciences, 17(10), 408–413. 10.1016/0968-0004(92)90010-7

da Veiga Leprevost, F., Haynes, S. E., Avtonomov, D. M., Chang, H.-Y., Shanmugam, A. K., Mellacheruvu, D., Kong, A. T., & Nesvizhskii, A. I. (2020). Philosopher: A versatile toolkit for shotgun proteomics data analysis. Nature Methods, 17(9), 869–870. 10.1038/s41592-020-0912-y

Demichev, V., Messner, C. B., Vernardis, S. I., Lilley, K. S., & Ralser, M. (2020). DIA-NN: Neural networks and interference correction enable deep proteome coverage in high throughput. Nature Methods, 17(1), 41–44. 10.1038/s41592-019-0638-x

Dengis, P. B., Nélissen, L. R., & Rouxhet, P. G. (1995). Mechanisms of yeast flocculation: Comparison of top-and bottom-fermenting strains. Applied and Environmental Microbiology, 61(2), 718–728. 10.1128/aem.61.2.718-728.1995

Deutsch, E. W., Bandeira, N., Perez-Riverol, Y., Sharma, V., Carver, J. J., Mendoza, L., Kundu, D. J., Wang, S., Bandla, C., Kamatchinathan, S., Hewapathirana, S., Pullman, B. S., Wertz, J., Sun, Z., Kawano, S., Okuda, S., Watanabe, Y., MacLean, B., MacCoss, M. J.,…Vizcaíno, J. A. (2023). The ProteomeXchange consortium at 10 years: 2023 update. Nucleic Acids Research, 51(D1), D1539–D1548. 10.1093/nar/gkac1040

D’Hautcourt, O., & Smart, K. A. (1999). Measurement of Brewing Yeast Flocculation. Journal of the American Society of Brewing Chemists, 57(4), 123–128. 10.1094/ASBCJ-57-0123

Djomehri, S. I., Gonzalez, M. E., da Veiga Leprevost, F., Tekula, S. R., Chang, H.-Y., White, M. J., Cimino-Mathews, A., Burman, B., Basrur, V., Argani, P., Nesvizhskii, A. I., & Kleer, C. G. (2020). Quantitative proteomic landscape of metaplastic breast carcinoma pathological subtypes and their relationship to triple-negative tumors. Nature Communications, 11(1), 1723. 10.1038/s41467-020-15283-z

Engler, C., Kandzia, R., & Marillonnet, S. (2008). A One Pot, One Step, Precision Cloning Method with High Throughput Capability. PLOS ONE, 3(11), e3647. 10.1371/journal.pone.0003647

European Organization For Nuclear Research & OpenAIRE. (2013). Zenodo. CERN. 10.25495/7GXK-RD71

Fasolo, J., Sboner, A., Sun, M. G. F., Yu, H., Chen, R., Sharon, D., Kim, P. M., Gerstein, M., & Snyder, M. (2011). Diverse protein kinase interactions identified by protein microarrays reveal novel connections between cellular processes. Genes & Development, 25(7), 767–778. 10.1101/gad.1998811

Gallone, B., Mertens, S., Gordon, J. L., Maere, S., Verstrepen, K. J., & Steensels, J. (2018). Origins, evolution, domestication and diversity of *Saccharomyces* beer yeasts. *Current Opinion in Biotechnology*, Food Biotechnology • Plant Biotechnology, 49, 148–155. 10.1016/j.copbio.2017.08.005

García-Gimeno, M. A., Muñoz, I., Ariño, J., & Sanz, P. (2003). Molecular characterization of Ypi1, a novel Saccharomyces cerevisiae type 1 protein phosphatase inhibitor. The Journal of Biological Chemistry, 278(48), 47744–47752. 10.1074/jbc.M306157200

Garnier, Simon, Ross, Noam, Rudis, Robert, Camargo, Pedro, A., Sciaini, Marco, Scherer, & Cédric. (2023). viridis(Lite)—Colorblind-Friendly Color Maps for R. 10.5281/zenodo.4678327

Gibson, B. R., Graham, N. S., Boulton, C. A., Box, W. G., Lawrence, S. J., Linforth, R. S. T., May, S. T., & Smart, K. A. (2010). Differential Yeast Gene Transcription during Brewery Propagation. Journal of the American Society of Brewing Chemists, 68(1), 21–29. 10.1094/ASBCJ-2009-1123-01

Gibson, B. R., Storgårds, E., Krogerus, K., & Vidgren, V. (2013). Comparative physiology and fermentation performance of Saaz and Frohberg lager yeast strains and the parental species Saccharomyces eubayanus. Yeast, 30(7), 255–266. 10.1002/yea.2960

Gimeno, C. J., Ljungdahl, P. O., Styles, C. A., & Fink, G. R. (1992). Unipolar cell divisions in the yeast S. cerevisiae lead to filamentous growth: Regulation by starvation and RAS. Cell, 68(6), 1077–1090. 10.1016/0092-8674(92)90079-r

Gorter de Vries, A. R., de Groot, P. A., van den Broek, M., & Daran, J.-M. G. (2017). CRISPR-Cas9 mediated gene deletions in lager yeast Saccharomyces pastorianus. Microbial Cell Factories, 16(1), 222. 10.1186/s12934-017-0835-1

Harashima, T., Anderson, S., Yates, J. R., & Heitman, J. (2006). The kelch proteins Gpb1 and Gpb2 inhibit Ras activity via association with the yeast RasGAP neurofibromin homologs Ira1 and Ira2. Molecular Cell, 22(6), 819–830. 10.1016/j.molcel.2006.05.011

Hinnebusch, A. G., & Fink, G. R. (1983). Positive regulation in the general amino acid control of Saccharomyces cerevisiae. Proceedings of the National Academy of Sciences of the United States of America, 80(17), 5374–5378. 10.1073/pnas.80.17.5374

Huber, W., von Heydebreck, A., Sueltmann, H., Poustka, A., & Vingron, M. (2003). Parameter estimation for the calibration and variance stabilization of microarray data. Statistical Applications in Genetics and Molecular Biology, 2, Article3. 10.2202/1544-6115.1008

Huber, W., von Heydebreck, A., Sültmann, H., Poustka, A., & Vingron, M. (2002). Variance stabilization applied to microarray data calibration and to the quantification of differential expression. *Bioinformatics (Oxford*, England), 18 *Suppl 1*, S96–104. 10.1093/bioinformatics/18.suppl_1.s96

Hunter, T. (2007). The age of crosstalk: Phosphorylation, ubiquitination, and beyond. Molecular Cell, 28(5), 730–738. 10.1016/j.molcel.2007.11.019

Jadav, R., Weiland, F., Noordermeer, S. M., Carroll, T., Gao, Y., Wang, J., Zhou, H., Lamoliatte, F., Muñoz, I., Toth, R., Macartney, T., Brown, F., Hastie, C. J., Alabert, C., Attikum, H. van, Zenke, F., Masson, J.-Y., & Rouse, J. (2023). Chemo-phosphoproteomic profiling with ATR inhibitors berzosertib and gartisertib uncovers new biomarkers and DNA damage response regulators (p. 2023.04.03.535285). bioRxiv. 10.1101/2023.04.03.535285

Joshua, I. M., Lin, M., Mardjuki, A., Mazzola, A., & Höfken, T. (2023). A Protein-Protein Interaction Analysis Suggests a Wide Range of New Functions for the p21-Activated Kinase (PAK) Ste20. International Journal of Molecular Sciences, 24(21), 15916. 10.3390/ijms242115916

Käll, L., Canterbury, J. D., Weston, J., Noble, W. S., & MacCoss, M. J. (2007). Semi-supervised learning for peptide identification from shotgun proteomics datasets. Nature Methods, 4(11), 923–925. 10.1038/nmeth1113

Khanam, T., Muñoz, I., Weiland, F., Carroll, T., Morgan, M., Borsos, B. N., Pantazi, V., Slean, M., Novak, M., Toth, R., Appleton, P., Pankotai, T., Zhou, H., & Rouse, J. (2021). CDKL5 kinase controls transcription-coupled responses to DNA damage. The EMBO Journal, 40(23), e108271. 10.15252/embj.2021108271

Kim, J., & Rose, M. D. (2015). Stable Pseudohyphal Growth in Budding Yeast Induced by Synergism between Septin Defects and Altered MAP-kinase Signaling. PLoS Genetics, 11(12), e1005684. 10.1371/journal.pgen.1005684

Kobayashi, O., Hayashi, N., Kuroki, R., & Sone, H. (1998). Region of Flo1 Proteins Responsible for Sugar Recognition. Journal of Bacteriology, 180(24), 6503–6510. 10.1128/jb.180.24.6503-6510.1998

Kong, A. T., Leprevost, F. V., Avtonomov, D. M., Mellacheruvu, D., & Nesvizhskii, A. I. (2017). MSFragger: Ultrafast and comprehensive peptide identification in mass spectrometry–based proteomics. Nature Methods, 14(5), 513–520. 10.1038/nmeth.4256

Kremer, L. P. M. (2019). ggpointdensity: A Cross Between a 2D Density Plot and a Scatter Plot. https://CRAN.R-project.org/package=ggpointdensity

*Lager Market Size, Share | Industry Report, 2035*. (n.d.). Retrieved July 18, 2026, from https://www.industryresearch.biz/market-reports/lager-market-113699

Laxman, S., & Tu, B. P. (2011). Multiple TORC1-associated proteins regulate nitrogen starvation-dependent cellular differentiation in Saccharomyces cerevisiae. PloS One, 6(10), e26081. 10.1371/journal.pone.0026081

Lee, M. E., DeLoache, W. C., Cervantes, B., & Dueber, J. E. (2015). A Highly Characterized Yeast Toolkit for Modular, Multipart Assembly. ACS Synthetic Biology, 4(9), 975–986. 10.1021/sb500366v

Lengeler, K. B., Davidson, R. C., D’souza, C., Harashima, T., Shen, W. C., Wang, P., Pan, X., Waugh, M., & Heitman, J. (2000). Signal transduction cascades regulating fungal development and virulence. Microbiology and Molecular Biology Reviews: MMBR, 64(4), 746–785. 10.1128/MMBR.64.4.746-785.2000

Liu, H., Styles, C. A., & Fink, G. R. (1993). Elements of the yeast pheromone response pathway required for filamentous growth of diploids. Science, 262(5140), 1741–1744. 10.1126/science.8259520

Lo, W. S., & Dranginis, A. M. (1996). FLO11, a yeast gene related to the STA genes, encodes a novel cell surface flocculin. Journal of Bacteriology, 178(24), 7144–7151. 10.1128/jb.178.24.7144-7151.1996

Lodolo, E. J., Kock, J. L. F., Axcell, B. C., & Brooks, M. (2008). The yeast Saccharomyces cerevisiae-the main character in beer brewing. FEMS Yeast Research, 8(7), 1018–1036. 10.1111/j.1567-1364.2008.00433.x

Lorenz, M. C., & Heitman, J. (1997). Yeast pseudohyphal growth is regulated by GPA2, a G protein alpha homolog. The EMBO Journal, 16(23), 7008–7018. 10.1093/emboj/16.23.7008

Lu, Z., & Hunter, T. (2009). Degradation of Activated Protein Kinases by Ubiquitination. Annual Review of Biochemistry, 78, 435–475. 10.1146/annurev.biochem.013008.092711

Maeda, T., Takekawa, M., & Saito, H. (1995). Activation of yeast PBS2 MAPKK by MAPKKKs or by binding of an SH3-containing osmosensor. Science, 269(5223), 554–558. 10.1126/science.7624781

Magalhães, F., Vidgren, V., Ruohonen, L., & Gibson, B. (2016). Maltose and maltotriose utilisation by group I strains of the hybrid lager yeast Saccharomyces pastorianus. FEMS Yeast Research, 16(5), fow053. 10.1093/femsyr/fow053

Mjikane, S. (2025, February 4). Beer’s Global Economic Footprint. Oxford Economics. https://www.oxfordeconomics.com/resource/beers-global-economic-footprint-2025/

Mösch, H. U. (2000). Pseudohyphal development of Saccharomyces cerevisiae. Contributions to Microbiology, 5, 185–200. 10.1159/000060354

Mösch, H. U., & Fink, G. R. (1997). Dissection of filamentous growth by transposon mutagenesis in Saccharomyces cerevisiae. Genetics, 145(3), 671–684. 10.1093/genetics/145.3.671

Müller-Auffermann, K., Caldera, A., Jacob, F., & Hutzler, M. (2015). Characterization of Different Bottom Fermenting Saccharomyces pastorianus Brewing Yeast Strains. BrewingScience, 68(3/4), 46–57. 10.23763/BrSc15-01mueller-auffermann

Norman, K. L., Shively, C. A., De La Rocha, A. J., Mutlu, N., Basu, S., Cullen, P. J., & Kumar, A. (2018). Inositol polyphosphates regulate and predict yeast pseudohyphal growth phenotypes. PLoS Genetics, 14(6), e1007493. 10.1371/journal.pgen.1007493

Ogata, T. (2012). Nitrogen starvation induces expression of Lg-FLO1 and flocculation in bottom-fermenting yeast. Yeast, 29(11), 487–494. 10.1002/yea.2928

Ogata, T., Izumikawa, M., Kohno, K., & Shibata, K. (2008). Chromosomal location of Lg-FLO1 in bottom-fermenting yeast and the FLO5 locus of industrial yeast. Journal of Applied Microbiology, 105(4), 1186–1198. 10.1111/j.1365-2672.2008.03852.x

Okuda, S., Yoshizawa, A. C., Kobayashi, D., Takahashi, Y., Watanabe, Y., Moriya, Y., Hatano, A., Takami, T., Matsumoto, M., Araki, N., Tabata, T., Iwasaki, M., Sugiyama, N., Kodera, Y., Tanaka, S., Goto, S., Kawano, S., & Ishihama, Y. (2025). jPOST environment accelerates the reuse and reanalysis of public proteome mass spectrometry data. Nucleic Acids Research, 53(D1), D462–D467. 10.1093/nar/gkae1032

Olivella, R., Chiva, C., Serret, M., Mancera, D., Cozzuto, L., Hermoso, A., Borràs, E., Espadas, G., Morales, J., Pastor, O., Solé, A., Ponomarenko, J., & Sabidó, E. (2021). QCloud2: An Improved Cloud-based Quality-Control System for Mass-Spectrometry-based Proteomics Laboratories. Journal of Proteome Research, 20(4), 2010–2013. 10.1021/acs.jproteome.0c00853

Orlova, M., Kanter, E., Krakovich, D., & Kuchin, S. (2006). Nitrogen availability and TOR regulate the Snf1 protein kinase in Saccharomyces cerevisiae. Eukaryotic Cell, 5(11), 1831–1837. 10.1128/EC.00110-06

Pan, X., & Heitman, J. (2000). Sok2 regulates yeast pseudohyphal differentiation via a transcription factor cascade that regulates cell-cell adhesion. Molecular and Cellular Biology, 20(22), 8364–8372. 10.1128/MCB.20.22.8364-8372.2000

Phipson, B., Lee, S., Majewski, I. J., Alexander, W. S., & Smyth, G. K. (2016). ROBUST HYPERPARAMETER ESTIMATION PROTECTS AGAINST HYPERVARIABLE GENES AND IMPROVES POWER TO DETECT DIFFERENTIAL EXPRESSION. The Annals of Applied Statistics, 10(2), 946–963. 10.1214/16-AOAS920

Pöhlmann, J., & Fleig, U. (2010). Asp1, a conserved 1/3 inositol polyphosphate kinase, regulates the dimorphic switch in Schizosaccharomyces pombe. Molecular and Cellular Biology, 30(18), 4535–4547. 10.1128/MCB.00472-10

Powell, C. D., Quain, D. E., & Smart, K. A. (2003). The impact of brewing yeast cell age on fermentation performance, attenuation and flocculation. FEMS Yeast Research, 3(2), 149–157. 10.1016/S1567-1356(03)00002-3

R Core Team. (2023). R: A Language and Environment for Statistical Computing. R Foundation for Statistical Computing. https://www.R-project.org/

Ram, K., & Wickham, H. (2023). wesanderson: A Wes Anderson Palette Generator. 10.32614/CRAN.package.wesanderson

Ritchie, M. E., Phipson, B., Wu, D., Hu, Y., Law, C. W., Shi, W., & Smyth, G. K. (2015). Limma powers differential expression analyses for RNA-sequencing and microarray studies. Nucleic Acids Research, 43(7), e47. 10.1093/nar/gkv007

Ryan, O., Shapiro, R. S., Kurat, C. F., Mayhew, D., Baryshnikova, A., Chin, B., Lin, Z.-Y., Cox, M. J., Vizeacoumar, F., Cheung, D., Bahr, S., Tsui, K., Tebbji, F., Sellam, A., Istel, F., Schwarzmüller, T., Reynolds, T. B., Kuchler, K., Gifford, D. K.,…Boone, C. (2012). Global gene deletion analysis exploring yeast filamentous growth. Science, 337(6100), 1353–1356. 10.1126/science.1224339

Salazar, A. N., Gorter de Vries, A. R., van den Broek, M., Brouwers, N., de la Torre Cortès, P., Kuijpers, N. G. A., Daran, J.-M. G., & Abeel, T. (2019). Chromosome level assembly and comparative genome analysis confirm lager-brewing yeasts originated from a single hybridization. BMC Genomics, 20(1), 916. 10.1186/s12864-019-6263-3

Sato, M., Maeba, H., Watari, J., & Takashio, M. (2002). Analysis of an inactivated Lg-*FLO1* gene present in bottom-fermenting yeast. Journal of Bioscience and Bioengineering, 93(4), 395–398. 10.1016/S1389-1723(02)80073-1

Sato, M., Watari, J., & Shinotsuka, K. (2001). Genetic Instability in Flocculation of Bottom-Fermenting Yeast. Journal of the American Society of Brewing Chemists, 59(3), 130–134. 10.1094/ASBCJ-59-0130

Seibel, K., Schmalhaus, R., Haensel, M., & Weiland, F. (2021). Molecular basis and regulation of flocculation in Saccharomyces cerevisiae and Saccharomyces pastorianus – a review. BrewingScience, 74(3/4), 39–50. 10.23763/BrSc21-03seibel

Shively, C. A., Eckwahl, M. J., Dobry, C. J., Mellacheruvu, D., Nesvizhskii, A., & Kumar, A. (2013). Genetic networks inducing invasive growth in Saccharomyces cerevisiae identified through systematic genome-wide overexpression. Genetics, 193(4), 1297–1310. 10.1534/genetics.112.147876

Shteynberg, D. D., Deutsch, E. W., Campbell, D. S., Hoopmann, M. R., Kusebauch, U., Lee, D., Mendoza, L., Midha, M. K., Sun, Z., Whetton, A. D., & Moritz, R. L. (2019). PTMProphet: Fast and Accurate Mass Modification Localization for the Trans-Proteomic Pipeline. Journal of Proteome Research, 18(12), 4262–4272. 10.1021/acs.jproteome.9b00205

Slowikowski, K. (2024). ggrepel: Automatically Position Non-Overlapping Text Labels with “ggplot2.” https://CRAN.R-project.org/package=ggrepel

Soares, E. V. (2011). Flocculation in Saccharomyces cerevisiae: A review. Journal of Applied Microbiology, 110(1), 1–18. 10.1111/j.1365-2672.2010.04897.x

Stewart, G. G. (2016). Saccharomyces species in the Production of Beer. Beverages, 2(4), 34. 10.3390/beverages2040034

Stratford, M., & Assinder, S. (1991). Yeast flocculation: Flo1 and NewFlo phenotypes and receptor structure. Yeast, 7(6), 559–574. 10.1002/yea.320070604

Teo, G. C., Polasky, D. A., Yu, F., & Nesvizhskii, A. I. (2021). Fast Deisotoping Algorithm and Its Implementation in the MSFragger Search Engine. Journal of Proteome Research, 20(1), 498–505. 10.1021/acs.jproteome.0c00544

Tyanova, S., Temu, T., & Cox, J. (2016). The MaxQuant computational platform for mass spectrometry-based shotgun proteomics. Nature Protocols, 11(12), 2301–2319. 10.1038/nprot.2016.136

Verstrepen, K. J., Derdelinckx, G., Verachtert, H., & Delvaux, F. R. (2003). Yeast flocculation: What brewers should know. Applied Microbiology and Biotechnology, 61(3), 197–205. 10.1007/s00253-002-1200-8

Walsh, E. P., Lamont, D. J., Beattie, K. A., & Stark, M. J. R. (2002). Novel Interactions of Saccharomyces cerevisiae Type 1 Protein Phosphatase Identified by Single-Step Affinity Purification and Mass Spectrometry. Biochemistry, 41(7), 2409–2420. 10.1021/bi015815e

Ward, M. P., Gimeno, C. J., Fink, G. R., & Garrett, S. (1995). SOK2 may regulate cyclic AMP-dependent protein kinase-stimulated growth and pseudohyphal development by repressing transcription. Molecular and Cellular Biology, 15(12), 6854–6863. 10.1128/MCB.15.12.6854

Wendland, J. (2014). Lager yeast comes of age. Eukaryotic Cell, 13(10), 1256–1265. 10.1128/EC.00134-14

Wickham, H. (2007). Reshaping Data with the reshape Package. Journal of Statistical Software, 21(12), 1–20.

Wickham, H. (2011). The Split-Apply-Combine Strategy for Data Analysis. Journal of Statistical Software, 40(1), 1–29.

Wickham, H. (2016). ggplot2: Elegant Graphics for Data Analysis. Springer-Verlag New York. https://ggplot2.tidyverse.org

Wickham, H. (2023). stringr: Simple, Consistent Wrappers for Common String Operations. https://CRAN.R-project.org/package=stringr

Wickham, H., Pedersen, T. L., & Seidel, D. (2023). scales: Scale Functions for Visualization. https://CRAN.R-project.org/package=scales

Wilson, W. A., & Roach, P. J. (2002). Nutrient-regulated protein kinases in budding yeast. Cell, 111(2), 155–158. 10.1016/s0092-8674(02)01043-7

Yang, K. L., Yu, F., Teo, G. C., Li, K., Demichev, V., Ralser, M., & Nesvizhskii, A. I. (2023). MSBooster: Improving peptide identification rates using deep learning-based features. Nature Communications, 14(1), 4539. 10.1038/s41467-023-40129-9

Zecha, J., Satpathy, S., Kanashova, T., Avanessian, S. C., Kane, M. H., Clauser, K. R., Mertins, P., Carr, S. A., & Kuster, B. (2019). TMT Labeling for the Masses: A Robust and Cost-efficient, In-solution Labeling Approach. Molecular & Cellular Proteomics0: MCP, 18(7), 1468–1478. 10.1074/mcp.TIR119.001385

