## Supplementary figures and images for "The hybrid yeast *Saccharomyces pastorianus* TUM 34/70 repurposes the pseudohyphal growth signaling network to establish a flocculation phenotype"

### MTHY8G_12_pYTK-STE-3_3.png

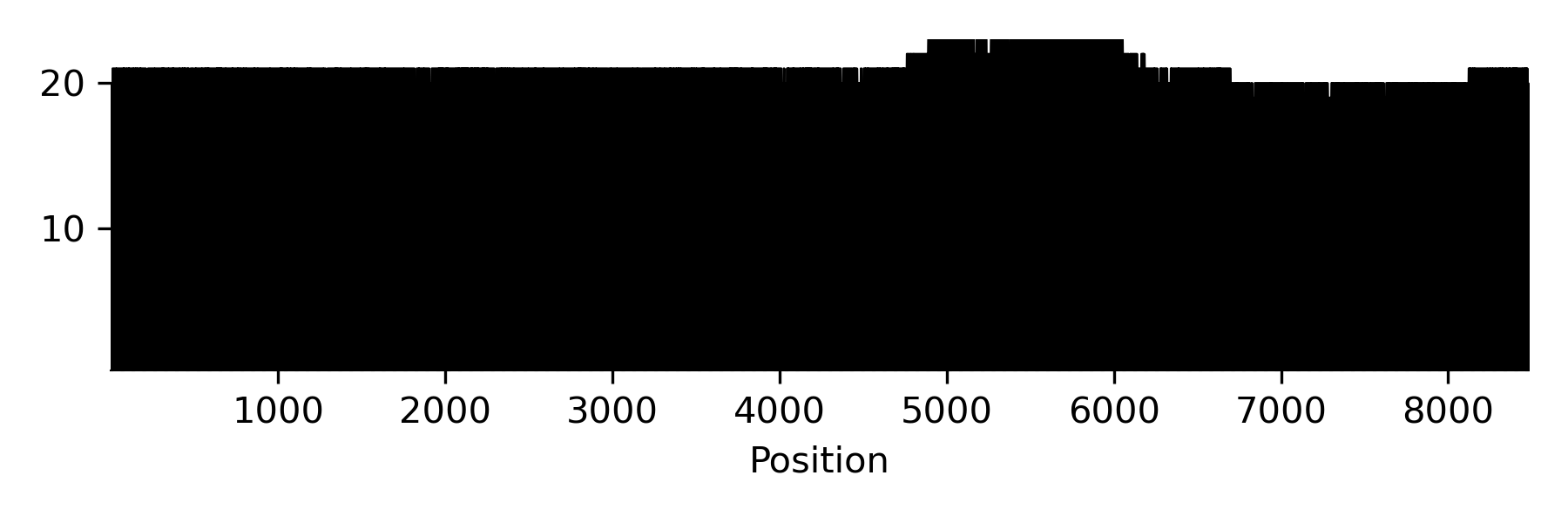

### MTHY8G_12_pYTK-STE-3_3.png

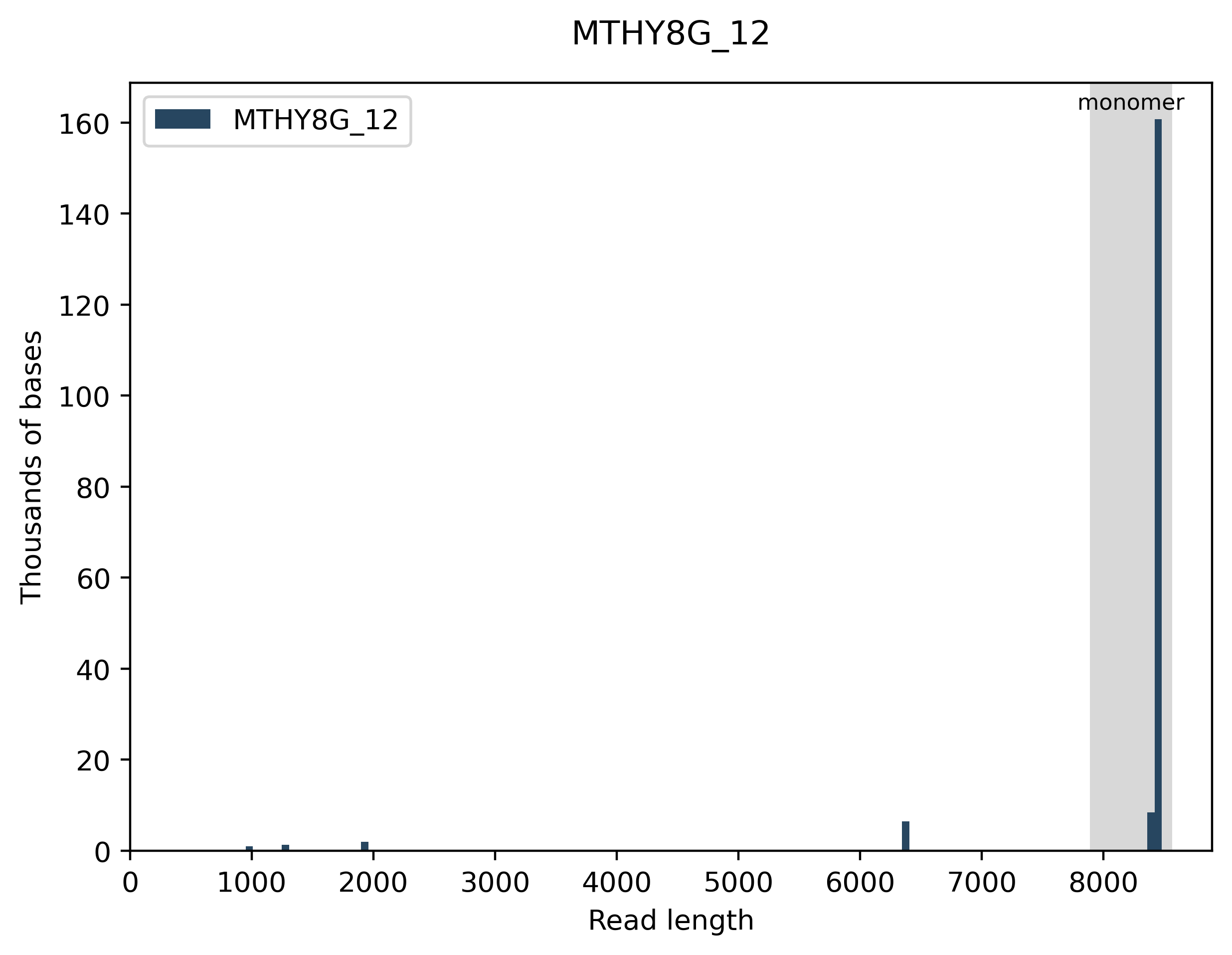

### T3P6XF_1_rerun_pYTK-GFP1-DO_1.png

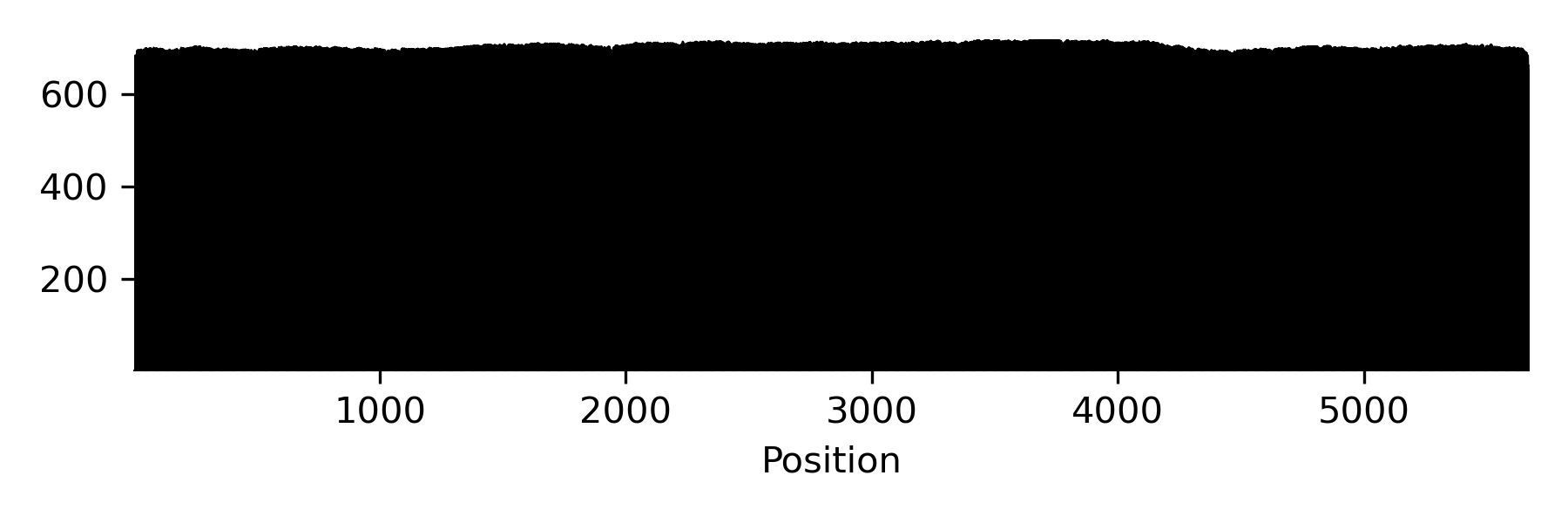

### T3P6XF_1_rerun_pYTK-GFP1-DO_1.png

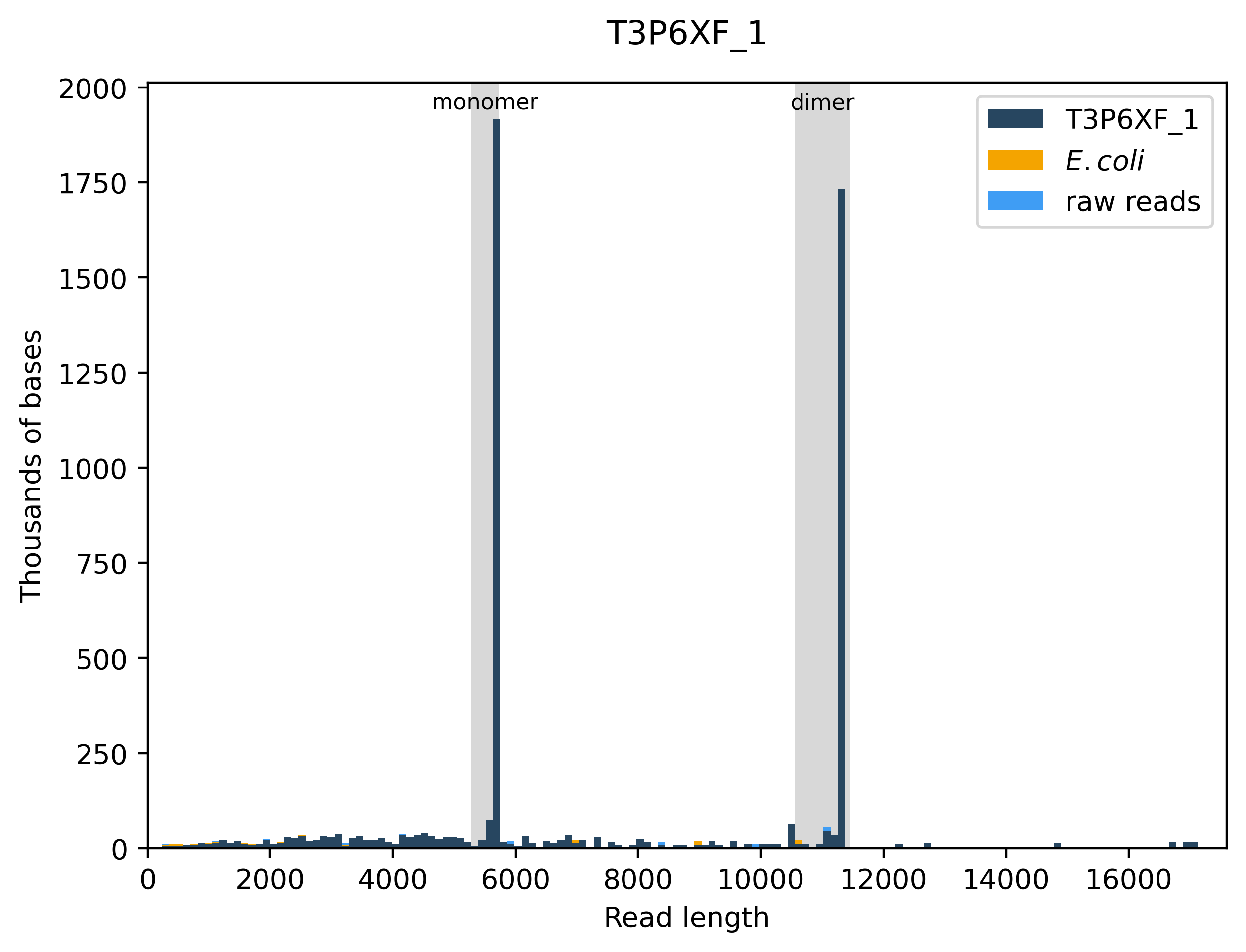

### ZDXKP2_1_pYTK-YPI_3__1.png

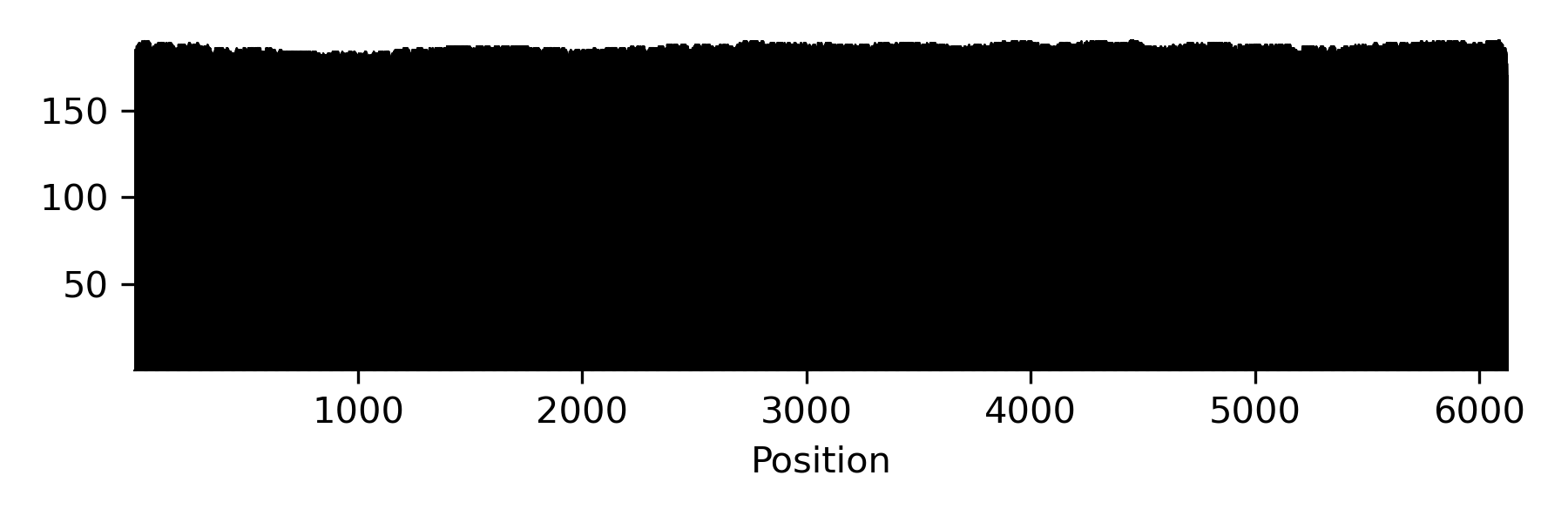

### ZDXKP2_1_pYTK-YPI_3__1.png

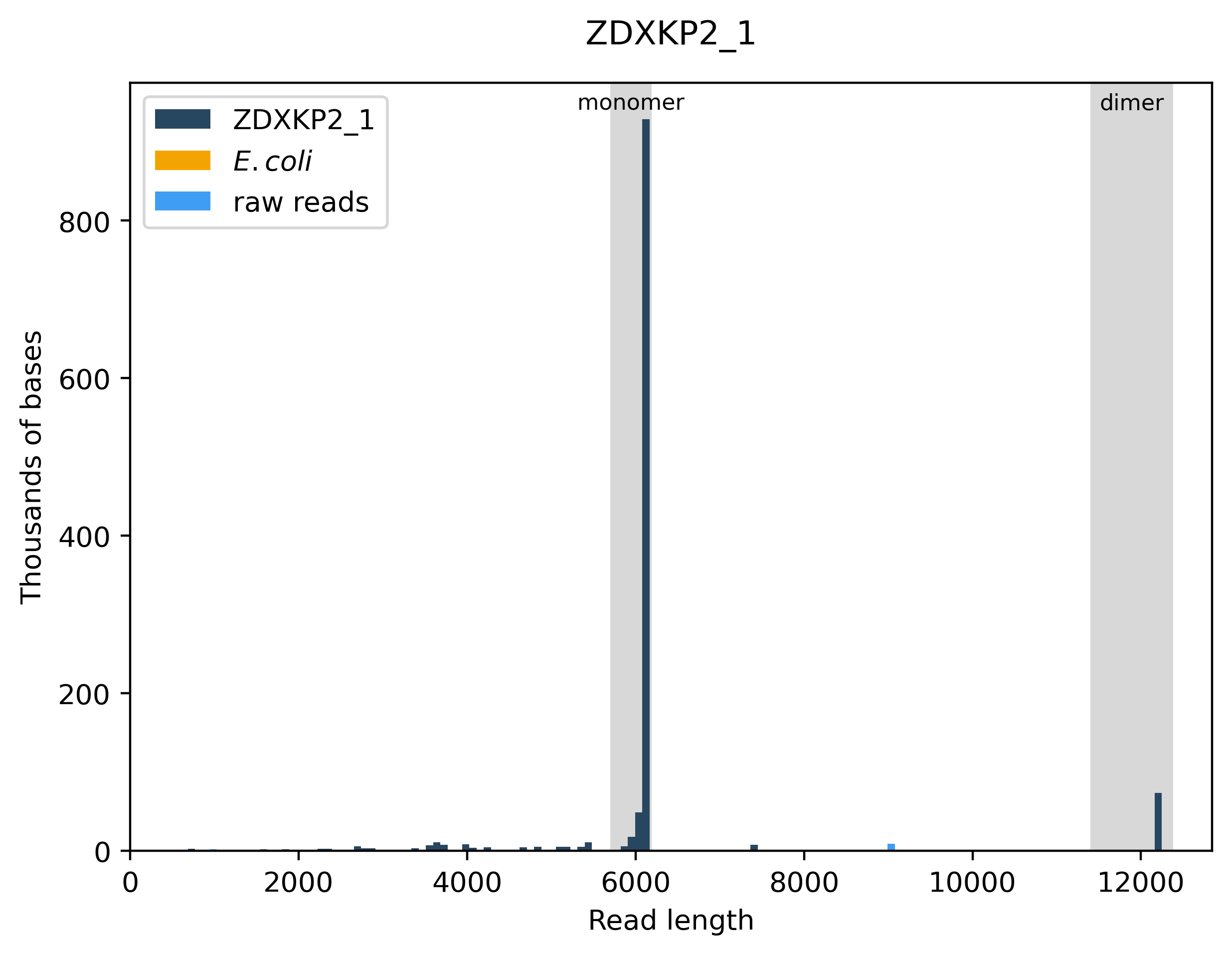
