## Supplementary_Data_and_Tables_Legends for "The hybrid yeast *Saccharomyces pastorianus* TUM 34/70 repurposes the pseudohyphal growth signaling network to establish a flocculation phenotype"

**Supplementary Data 1 – Sequencing results for validation of the used plasmids.**

**Supplementary Table 1 – List of used primers.**

**Supplementary Table 2 – Proteomics results.** Columns 10-43 depict the VSN transformed TMT reporter intensities. Positive logFC indicates a higher abundant peptide in the first condition mentioned under the “Contrast” column. Non-Floc: Yeast showing overall non-flocculent phenotype, Sed.: Sedimented subpopulation of yeast after 15 min rest in overall flocculent sample, Non-Sed.: Non-sedimented subpopulation of yeast after 15 min rest in overall flocculent sample. S: Samples supplemented with glutamine & glutamic acid, NS: Non-supplemented samples. Double-digit numbers indicate proteomics sampling time point after inoculation, last number indicates the (biological) replicate.

**Supplementary Table 3 – Phosphoproteomics results *STE20* and *YPI1* overexpression.** Modified.Sequence depicts the amino acid sequence with the modified amino acid mass in square brackets (S[167]: phospho-serine, T[181]: phospho-threonine, Y[243]: phospho-tyrosine, M[147]: oxidation of methionine, C[160]: carbamidomethylation of cysteine, K[432]: TMTpro modified lysine. Column “Qvalue “indicates the FDR of the peptide identification, “Protein.Start” and “Protein.End” localize the peptide within the amino acid sequence of the protein, “Assigned.Modifications” indicate the modifications with the corresponding amino acid position in the identified peptide sequence. “STY.79.9663 “ places the assigned modifications within the identified amino acid sequence and gives the localization probability in brackets. ” STY.79.9663.Best.Localization” indicates the value of the most probable localizations. Columns containing “WT” followed by 1-5 indicates the VSN transformed TMT reporter ion intensity of the wild-type yeast and the number indicates the biological replicate, “CTRL” of the 3 Ala control and STE20/YPI1 of the respective overexpressed gene.
