## Supplementary_Figures for "The hybrid yeast *Saccharomyces pastorianus* TUM 34/70 repurposes the pseudohyphal growth signaling network to establish a flocculation phenotype"

### Slide 1
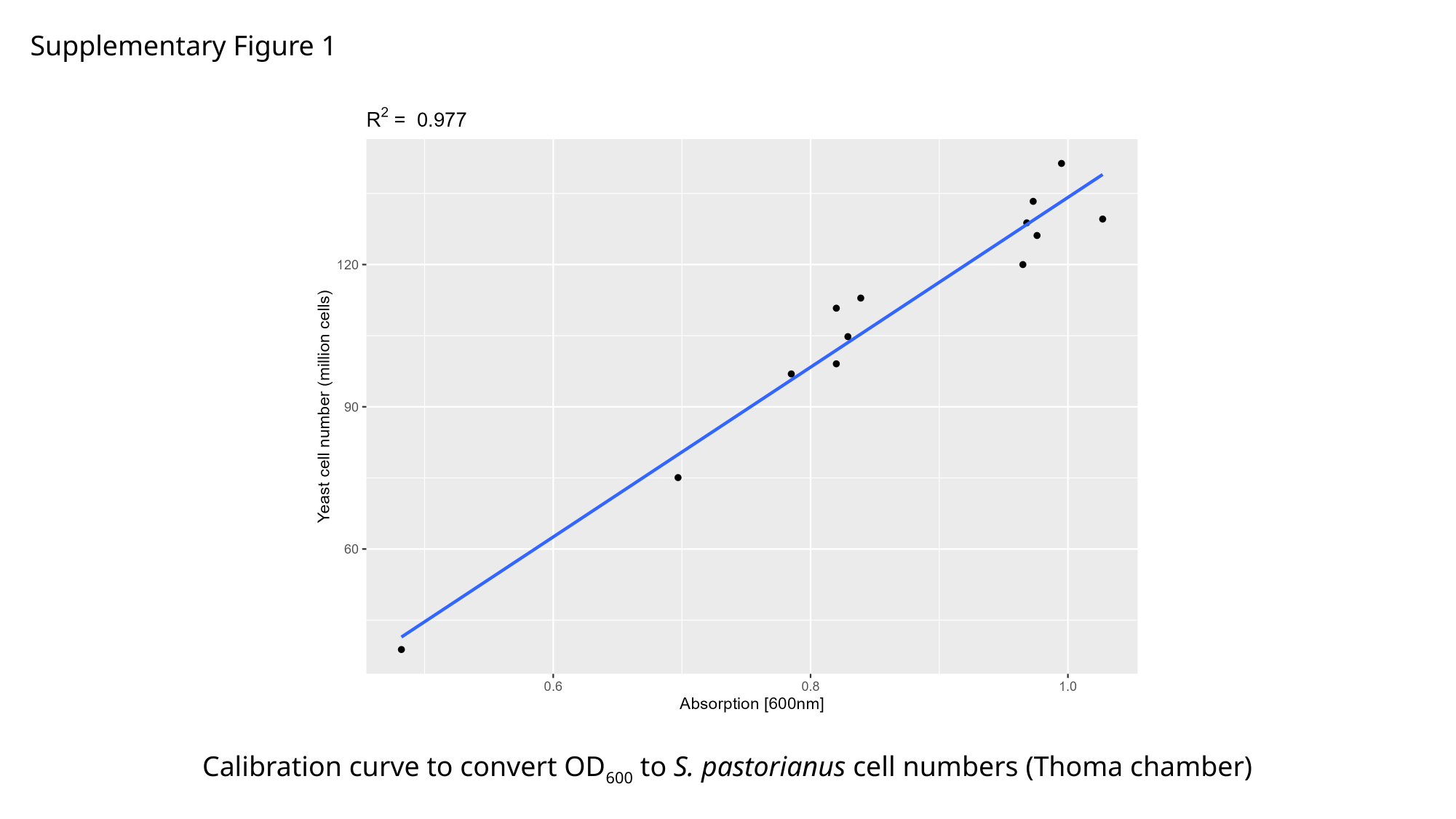

Supplementary Figure 1
Calibration curve to convert OD_600_ to S. pastorianus cell numbers (Thoma chamber)

### Slide 2
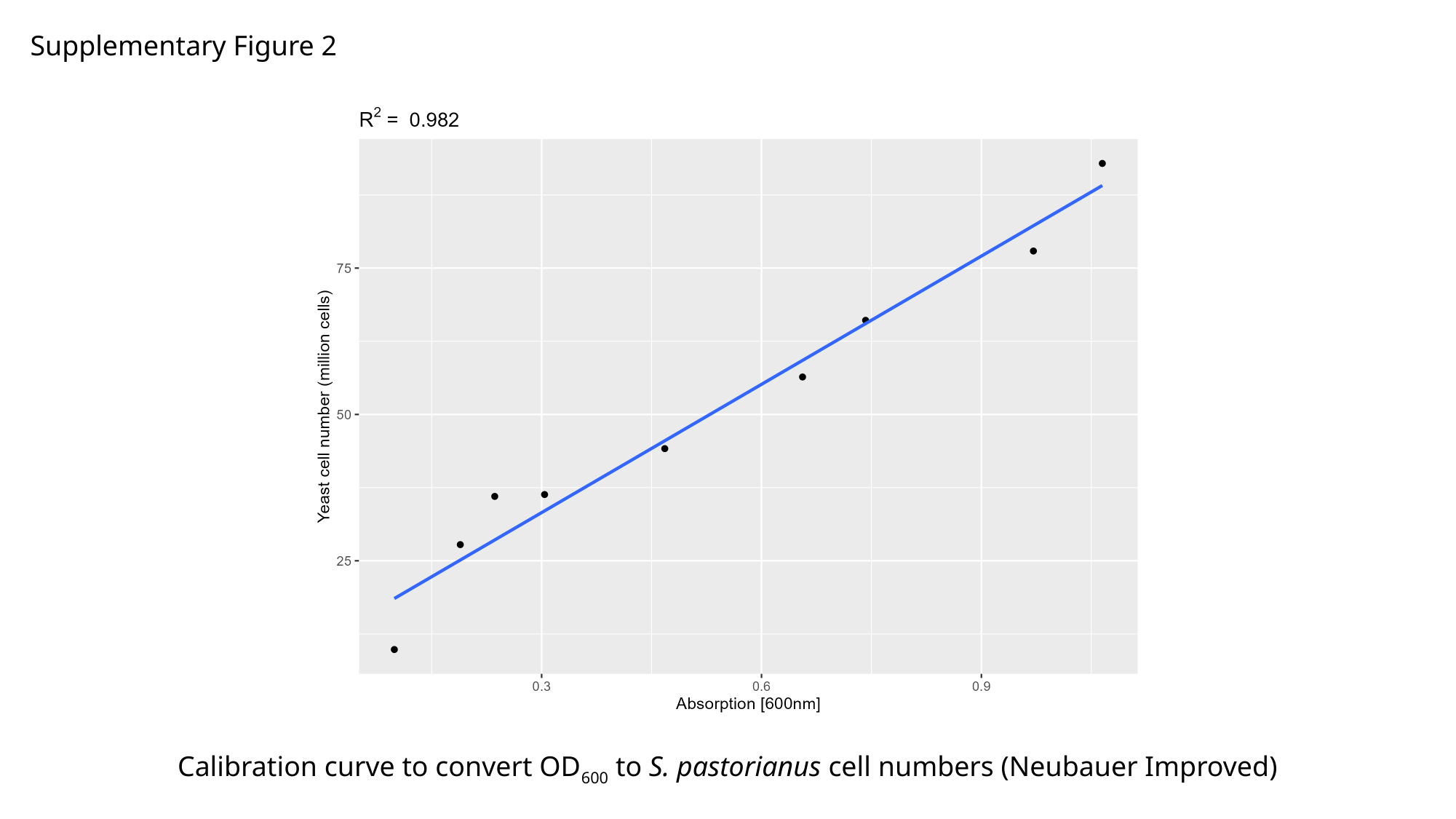

Supplementary Figure 2
Calibration curve to convert OD_600_ to S. pastorianus cell numbers (Neubauer Improved)

### Slide 3
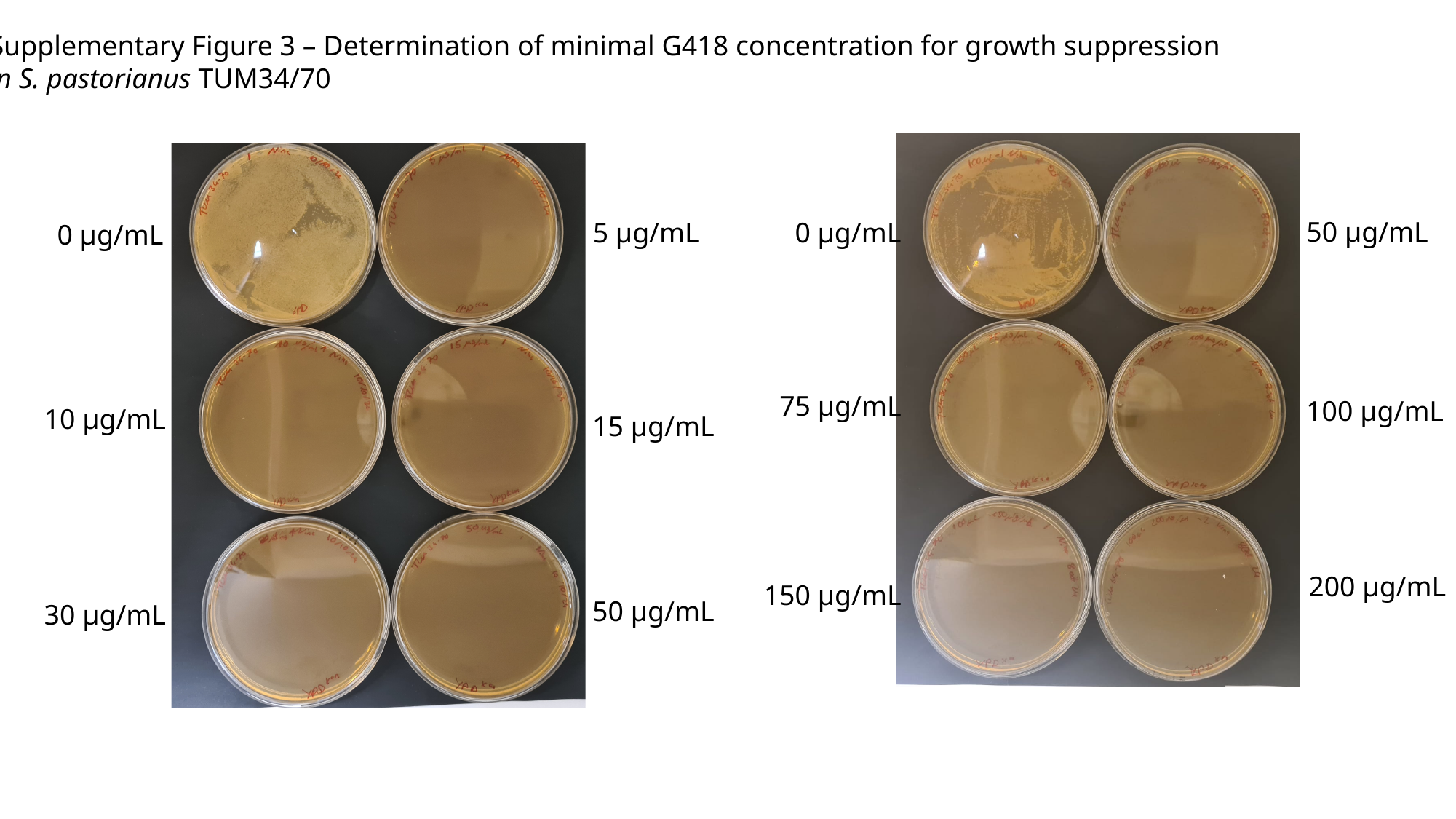

Supplementary Figure 3 – Determination of minimal G418 concentration for growth suppression
in S. pastorianus TUM34/70
50 µg/mL
5 µg/mL
0 µg/mL
0 µg/mL
75 µg/mL
100 µg/mL
10 µg/mL
15 µg/mL
200 µg/mL
150 µg/mL
50 µg/mL
30 µg/mL
